# Tabular foundation model predicts alternative lengthening of telomeres (ALT) and identifies SMARCAL1 as a target in ALT-driven cancers

**DOI:** 10.64898/2026.03.04.709438

**Authors:** D Bennett, M Wierdl, S Chakraborty, JD Johnson, V Akingbehin, D Estevez-Prado, Y Feng, JJ Morsby, J Harper, MA Mohammad Nezhady, M Pan, N Ocasio-Martinez, JL Catlett, TB Nations, NV Dharia, A Robichaud, A Herman, X Zhang, J Kelly, H Inam, S Stokes, J Steele, M Rusch, K Wienand, CD Campbell, G Alexe, F Vazquez, G Getz, CWM Roberts, CG Mullighan, EA Sweet-Cordero, CP Reynolds, B Koneru, AA Shelat, AD Durbin, E Bernstein, X Ma, K Stegmaier, P Geeleher, LM Guenther

## Abstract

Alternative lengthening of telomeres (ALT) is a telomerase-independent pathway used by aggressive cancers to maintain their replicative immortality. Because ALT is absent from normal human cells, it is an appealing target for cancer therapy, but the lack of ability to determine ALT status at scale has hindered discovery. Here, we developed ALTitude, a machine learning method from a tabular foundation model that infers ALT from cell line whole genome sequencing data, without need for paired germline analysis. We deployed ALTitude across the DepMap, doubling the number of known ALT+ cancer models. Systematic integration of ALTitude with CRISPR-Cas9 screens yielded the selective dependency on *SMARCAL1* in ALT+ cell lines, where we show it stabilizes the ALT phenotype. Acute depletion of *SMARCAL1* leads to G2/M arrest, mitotic catastrophe, and cell death. These data provide a valuable resource for studying ALT-related genomic features and present *SMARCAL1* as a therapeutic target for ALT+ malignancies.

---

Telomeres are repetitive nucleotide sequences capping the ends of eukaryotic chromosomes. They play an essential role in maintaining genomic stability and preventing chromosome degradation. Shortening of telomeres, which occurs with each healthy cell division, gradually leads to chromosomal instability and subsequent cell death.^1^ Replicative immortality, recognized as a hallmark of cancer cells, cannot be achieved without maintenance of telomere length.^2^ Most human cancers achieve this through constitutive activation of the enzyme telomerase. A minority of tumors rely instead on a telomerase-independent telomere maintenance mechanism (TMM) known as Alternative Lengthening of Telomeres (ALT),^3^ however, experimental methodology leads to different estimations of the prevalence of this mechanism. While 10-15% of tumors are found to lack telomerase activity, several recent studies investigating ALT using multiple assays in human tumor samples found the prevalence to be 2.5-5%, with wide variation between tumor subtypes.^4,5^ The highest reported subtypes that are characterized by ALT include sarcomas and high-grade brain tumors, with the lowest being epithelial tumors.^6,7^ ALT cells are characterized by heterogeneous telomere lengths, elevated levels of extrachromosomal telomeric DNA, and homologous recombination at telomeric loci.^8,9^ Generally, patients with ALT+ tumors fare poorly, with lower overall survival than their telomerase positive counterparts.^6,10–12^ Importantly, ALT is not found in normal, non-cancerous human cells.^13^ Therefore, targeting ALT in cancer could have a wide therapeutic window.

Detecting ALT has traditionally relied on experimental techniques such as C-circle assays to measure quantity of extrachromosomal telomeric DNA circles, telomere-specific fluorescence in situ hybridization (FISH) to identify ultrabright nuclear telomeric foci, or electron microscopy to detect recombination intermediates.^6,14^ Though sensitive, especially when combining orthogonal evidence across multiple assays, integrating these methods is labor-intensive, requires specialized reagents, and hinges on subjective judgment; thus, it is not easily applied at scale. As large-scale cancer ‘omics’ datasets become increasingly available, there have been recent efforts to infer ALT using computational approaches. With whole-genome sequencing (WGS) data, it is now possible to derive detailed telomere features, such as coverage patterns, structural variants, and the abundance of telomere-specific motifs, which may signal the presence or absence of ALT in a tumor.^15–18^ However, such methods have historically relied on paired germline sequencing and can suffer from sequencing batch effects. This hinders their application in diverse cohorts of preclinical cancer models, where germline information is not available, and/or WGS has typically been performed using varying technologies. Other efforts have sought to work around this by using proteomics and transcriptional data to computationally derive ALT and telomerase activity signatures,^5^ though these approaches have challenges including high false positive rates. We reasoned that to scale ALT calls for broader preclinical studies, as well as potential future applications, such as biomarker stratification in clinical trials, a streamlined, inexpensive methodology, retaining ultrahigh precision, is needed.

Here, we report a tabular foundation-based model (we call ‘ALTitude’) for predicting ALT in cancer cell lines based on WGS data, which we show achieves near perfect accuracy. Critically, the method does not require paired germline sequencing data, and automatically detects and corrects for sequencing batch, making it suitable for application in diverse aggregated panels of cancer cell lines sequenced on different platforms and across different centers. We applied ALTitude to pediatric and adult cancer cell lines with available WGS in the Cancer Dependency Map database (DepMap: ALTitude calls are integrated into www.depmap.org).^19–21^ We then combined ALT predictions in DepMap with somatic mutational and structural variant data from WGS and RNA-sequencing (RNA-seq), as well as genome-scale loss-of-function (LOF) CRISPR screening to identify selective vulnerabilities specific to ALT-positive (ALT+) tumor populations. One gene, SWI/SNF-Related Matrix-Associated Actin-Dependent Regulator of Chromatin Subfamily A-Like Protein 1 (*SMARCAL1*) emerged as a strong dependency in many ALT+ cancers. We go on to show that, in a subset of ALT+ tumor lines that have *ATRX/DAXX* loss-of-function alterations, SMARCAL1 functions as a synthetic lethal partner, with these genes mechanistically and functionally compensating for each other’s loss. Furthermore, we find that SMARCAL1-induced replication stress occurs regardless of ALT status, but that ALT+ cells are incapable of resolving the subsequent DNA damage, leading to the selective fitness effect in the ALT context. Overall, this work introduces a new, easy-to-apply approach for investigating ALT at scale, which yielded the identification of a biomarker-informed potential drug target for some of the most difficult-to-treat cancers.

## Results

### ALTitude is a tabular foundation-based model that can reliably identify ALT+ cancer cell line models in DepMap

To systematically identify and exploit vulnerabilities in ALT+ cancer cell line models, we devised a methodology to determine ALT status from the genomics data collected in cell lines from the DepMap. We developed ALTitude, a machine learning framework that leverages multiple telomere-relevant features estimated from WGS data (**Fig. 1a**). To meaningfully integrate these features across sequencing batches, we first developed a principal component regression-based normalization approach that removes technical variation, which is estimated from genome-wide variation in sequencing depth (see **Methods**; **Extended Data Fig. 1**). We trained ALTitude using 82 adult and pediatric (20 ALT+ and 62 non-ALT) cancer cell lines with orthogonally validated ALT status determined by C-circle assays, ALT-associated PML bodies (APB) assays,^22^ telomeric FISH, and/or assessment of telomerase activity with TRAP assays, which we curated from the existing literature (**Extended Data Table 1; Extended Data Fig. 2**). The model leveraged five feature categories: *TERT* gene expression, telomere length estimates, frequencies of 49 telomeric variant repeats (6-mer permutations containing GGG), telomere fusion rates, and the prevalence of neo-telomere structures (see **Methods**). We performed PCA of these telomere features in our 82 training cell lines, which revealed clear separation of ALT+ and non-ALT samples along PC1 (52.5 % variance). The PC loadings showed that neo-telomere structures and telomere fusions were the strongest contributors to the ALT+ axis, while *TERT* expression loaded in the opposite direction, consistent with higher *TERT* expression in non-ALT samples (**Fig. 1b**).

**Fig. 1.**
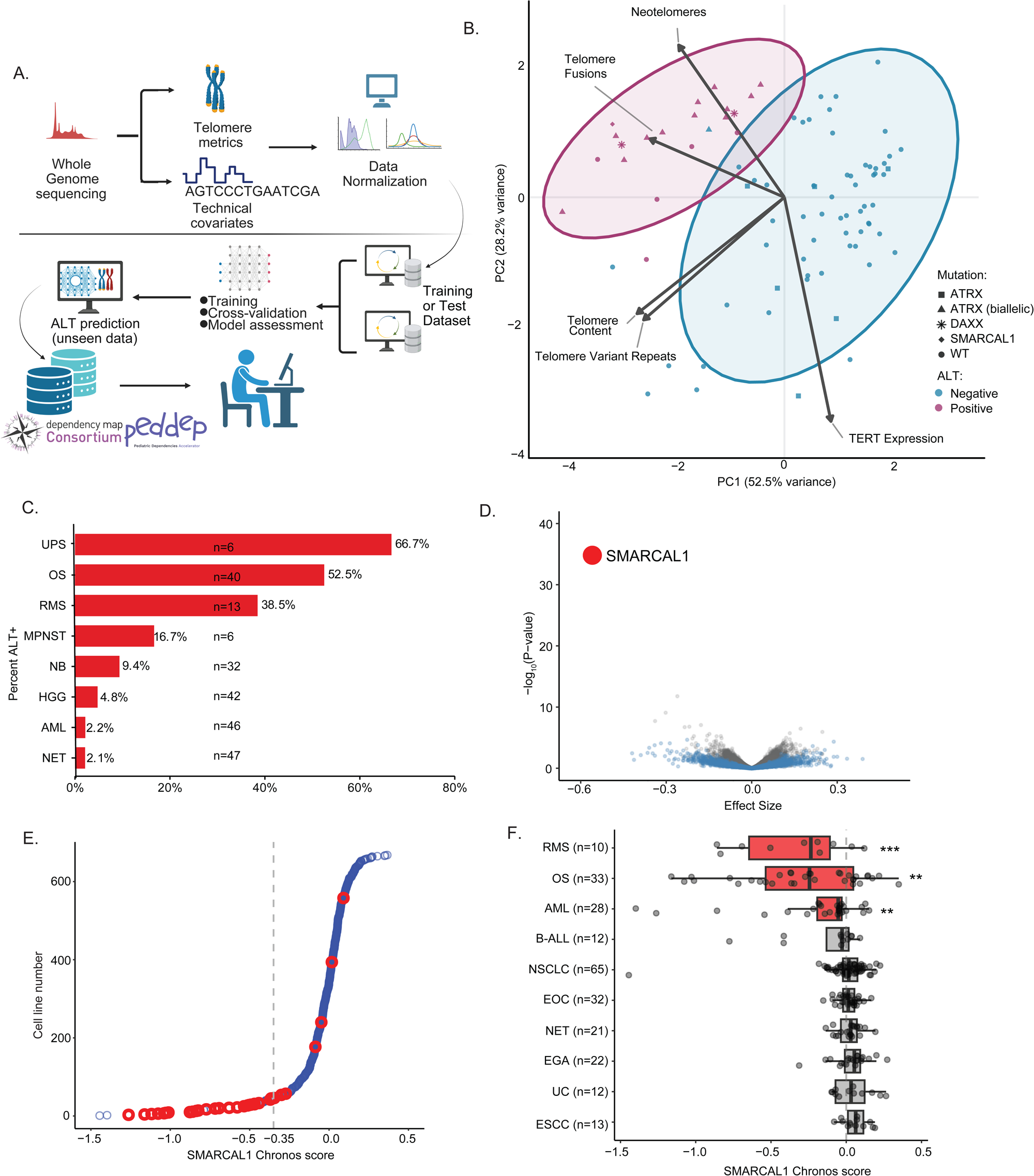
A whole genome sequencing-based machine learning-based classification of alternative lengthening of telomeres (ALT) identifies ALT-specific dependencies in cancer cell lines. a. Schematic overview of the computational workflow for ALT detection. Whole genome sequencing data undergoes extraction of telomere metrics (telomere content, C-circle annotation, telomere variant repeats) and technical covariates, followed by data normalization. A machine learning model incorporating attribute selection, training, cross-validation, and model assessment is applied to generate ALT predictions for unseen data. The model outputs feed into the DepMap web database and downstream drug discovery and validation pipelines. **b.** PCA of telomeric features characterizing ALT+ cell lines in the training dataset. Points represent individual cell lines colored by ALT assay status (blue = Negative, pink = Positive) with shapes indicating mutation status. Arrows show the top contributing features to each principal component. Ellipses indicate the 90% probability contour for each group assuming a bivariate normal distribution. PC1 and PC2 explain 52.5% and 28.2% of variance, respectively. **c.** Proportion of predicted ALTitude positive calls by disease type. Bars represent percent ALT models. Total sample sizes per disease indicated for each. **d.** Volcano plot showing differential dependency genes between ALT-positive and ALT-negative samples predicted by ALTitude. Effect size (x-axis) is plotted against statistical significance (-log_10_ p-value, y-axis). *SMARCAL1* is highlighted in red. e. Waterfall plot of ALT (red) vs. non-ALT (blue) cell lines as called by ALTitude. *SMARCAL1* Chronos score is shown on the x-axis. f. Bar and whisker plot demonstrating *SMARCAL1* Chronos score (x-axis) grouped by lineage, ordered on limma test significance of differential selectivity of Chronos scores within disease vs remaining diseases. Red represents enriched diseases. Top 10 diseases with > 5 cell line models are shown. ** p<0.01, *** p<0.001 by limma. UPS = Undifferentiated Pleomorphic Sarcoma; OS = osteosarcoma; RMS = rhabdomyosarcoma; MPNST = malignant peripheral nerve sheath tumor; NB = neuroblastoma; HGG = diffuse glioma; NET = lung neuroendocrine tumor; AML = acute myeloid leukemia; B-ALL = B-Acute Lymphoblastic Leukemia; NSCLC = non-small cell lung cancer; EOC = ovarian epithelial tumor; EGA = esophagogastric adenocarcinoma; UC = bladder uroepithelial carcinoma; ESCC = esophageal squamous cell carcinoma.

We evaluated the performance of nine different machine learning models: Random Forest, Logistic Regression, XGBoost, CatBoost, HistGradientBoosting, AdaBoost, LightGBM, ExtraTrees, and the neural network-based foundation model TabPFN2.^23–30^ Data were split into training (*N* = 57) and held-out sets (*N* = 25). Models were trained and evaluated using leave-one-out cross-validation (LOOCV) on the training set, where the performance ranged from AUROC of 0.975 (XGBoost) to 1.0 (LightGBM, CatBoost, HistGB, Logistic Regression & TabPFN2). On the held-out set all models performed with an AUC of 1 with excellent sensitivity (1.0) and specificity (0.947 - 1.0), demonstrating potential robust generalization to unseen samples (**Extended Data Table 2**). To further evaluate the relative robustness of the different machine learning models, we performed a feature ablation experiment across 20 feature configurations (**Extended Data Table 3**). TabPFN ranked first in both mean AUROC (0.961) and mean PR-AUC (0.909), indicating consistently strong performance across varying feature subsets (**Extended Data Fig. 3**), demonstrating superior robustness to feature perturbation. TabPFN is a neural network foundation model-based approach recently reported as state-of-the-art on tabular-scale data, and this is consistent with our evaluations.^30^ We thus selected TabPFN, fitting the final model on the entire (*N* = 82) cell line training set.

To validate ALTitude on independent data, we benchmarked against a recent cell line dataset with ALT status determined by C-circles, TRAP assays, and telomere content by qPCR (*Wu et al*.).^5^ Among Wu *et al*’s profiled cell lines for which WGS data was available from DepMap, ALTitude achieved a 0% false positive rate (0/440 lines mis-called ALT+). Two adenocarcinoma lines, SKLU1 and HS746T, classified as ALT by *Wu et al.* were called non-ALT by ALTitude. An additional 7 cell lines called non-ALT by ALTitude were annotated as having dual-positive (DP) TMM characteristics. In total, ALTitude predicted 10 cell lines concordantly ALT+ with Wu et al., and 440 concordantly non-ALT (**Extended Data Fig. 4**).

We finally compared ALTitude to a state-of-the-art RNA-based ALT classifier reported recently by *Wu et al.*, performing this analysis *using 5-fold cross-validation* on 82 cell lines with curated ALT calls overlapping between datasets.^5^ When implementing the published method in cross-validation (placing feature selection within each CV fold to avoid information leakage), the RNA model achieved AUROC of 0.946 compared to ALTitude’s AUROC of 1.0. ALTitude demonstrated superior specificity (100% vs 77%), and precision (100% vs 58.8%) (**Extended Data Fig. 4; Extended Data Table 4-5**), yielding far fewer false positive ALT calls. Our ALTitude results, which have been made publicly available in the DepMap web portal, provide the first comprehensive landscape of accurate ALT status predictions across a large panel of cancer cell line models from WGS data, meaning that ALT status can now be reliably determined prospectively for future cell line models.

### ALTitude predictions nominate novel ALT-specific vulnerabilities in DepMap

We next applied ALTitude to 906 DepMap cell lines for which WGS was available, predicting 19 new, previously undescribed ALT+ models, for a total of 4.3% (*N* = 39) ALT+ models across DepMap (**Extended Data Fig. 5, Extended Data Table 1**). This is similar to prevalence predicted across cancer patients using orthogonal methods, affirming the accuracy of our pipeline.^4^ We did not find any evidence in the literature to support the new ALT+ lines being telomerase dependent (e.g. reported by TRAP assay). As expected, the ALT cell lines were enriched in lineages corresponding to known ALT-prevalent cancers including undifferentiated pleomorphic sarcomas (UPS; n=6, 66.7%) and osteosarcoma (OS; n=40, 52.5%) (**Fig. 1c**).^4^

Next, to determine if our ALT predictions could be used to identify ALT-specific vulnerabilities, we jointly analyzed the ALTitude output with the genome-scale loss-of-function CRISPR screening data collected in DepMap, comparing the dependency scores for each of ∼20,000 genes between our 39 ALT+ and 867 non-ALT cell lines (see Methods). We identified *SMARCAL1* as the top differential dependency (*FDR* = 1.141 × 10^-31^, *β* = -0.561; **Extended Data Table 6**), with a greater fitness effect of CRISPR knockout (KO) in ALT+ cell lines (**Fig. 1d**). *SMARCAL1* has previously been described as a telomere regulator, including roles in maintenance of ALT+ and non-ALT telomeres, where it tempers replication stress.^31,32^ Recently, *SMARCAL1* was observed as a dependency in ALT+ high-grade gliomas, highlighting the accuracy and reproducibility of our ALTitude pipeline.^33^ In total, 45 DepMap cell lines had a moderate to strong dependency on *SMARCAL1* (defined as CHRONOS score of <-0.35, see **Methods**) and of these, 26 (57.8%) were called ALT+ by ALTitude (**Fig. 1e, Extended Data Table 1**). This suggests that there are a small number of ALT+ cell lines that are not *SMARCAL1*-dependent, and that *SMARCAL1* also represents a vulnerability in a small minority of non-ALT context(s), though the mechanisms are not known (see **Discussion**). We next examined the disease lineages enriched for dependency on *SMARCAL1* in ALT+ DepMap cell lines. Notably, rhabdomyosarcoma and osteosarcoma, both pediatric solid tumors, and acute myeloid leukemia (AML), specifically pediatric AML cases, were significantly more dependent on *SMARCAL1* than any other lineages (**Fig. 1f**).

### *SMARCAL1* is an ALT-associated dependency in osteosarcoma *in vitro* and confirmed in a pooled *in vivo* screen

We observed that, generally, DepMap pediatric cell lines showed greater enrichment for *SMARCAL1* dependency than adult cell lines, and that, as expected, most pediatric *SMARCAL1*-dependent lines were ALT+ (20 of 27, 74.1%) (**Fig. 2a-b**). Based on these patterns, we hypothesized that *SMARCAL1* represented a valuable new therapeutically relevant vulnerability in high-risk pediatric cancers where actionable drug targets are very limited. We first validated *SMARCAL1* dependency *in vitro* in osteosarcoma, a common pediatric tumor with high prevalence of ALT+ and strong *SMARCAL1* dependency in our dataset. Osteosarcoma is a genomically complex malignancy with many heterogeneous structural rearrangements and copy number changes, but few recurrent somatic mutations.^34,35^ Historic rates of ALT in osteosarcoma are among the highest of all cancers, estimated between 40-60%.^3,22,36^ Since there have been virtually no new treatment strategies for children with osteosarcoma in over forty years,^37^ finding a biomarker-defined target for ALT+ osteosarcoma would be transformative for this disease population. We performed lentiviral CRISPR KO in a panel of osteosarcoma cell lines included in the DepMap, demonstrating growth inhibition by CellTiter-Glo® (CTG) in ALT+ cell lines, with no significant viability changes in ALT negative osteosarcoma cell lines (**Fig. 2c-d**). To ensure broader application of this target beyond osteosarcoma, we confirmed viability effects of *SMARCAL1* KO in three ALT+ rhabdomyosarcoma cell lines by CTG (**Extended Data Fig. 6a-b**). As an orthogonal method to perturb and study SMARCAL1 function, we used the FKBP12^F36V^-degron system to generate an inducible protein degradation system (dTAG) on the c-terminus of *SMARCAL1* in the ALT+ osteosarcoma cell line G292.^38,39^ Degradation of FKBP12^F36V^-SMARCAL1 in these cells with CRISPR KO of endogenous *SMARCAL1* significantly reduced viability by CTG, as well as anchorage-independent growth in methylcellulose assays compared to controls (**Extended Data Fig. 6c-f**).

**Fig. 2.**
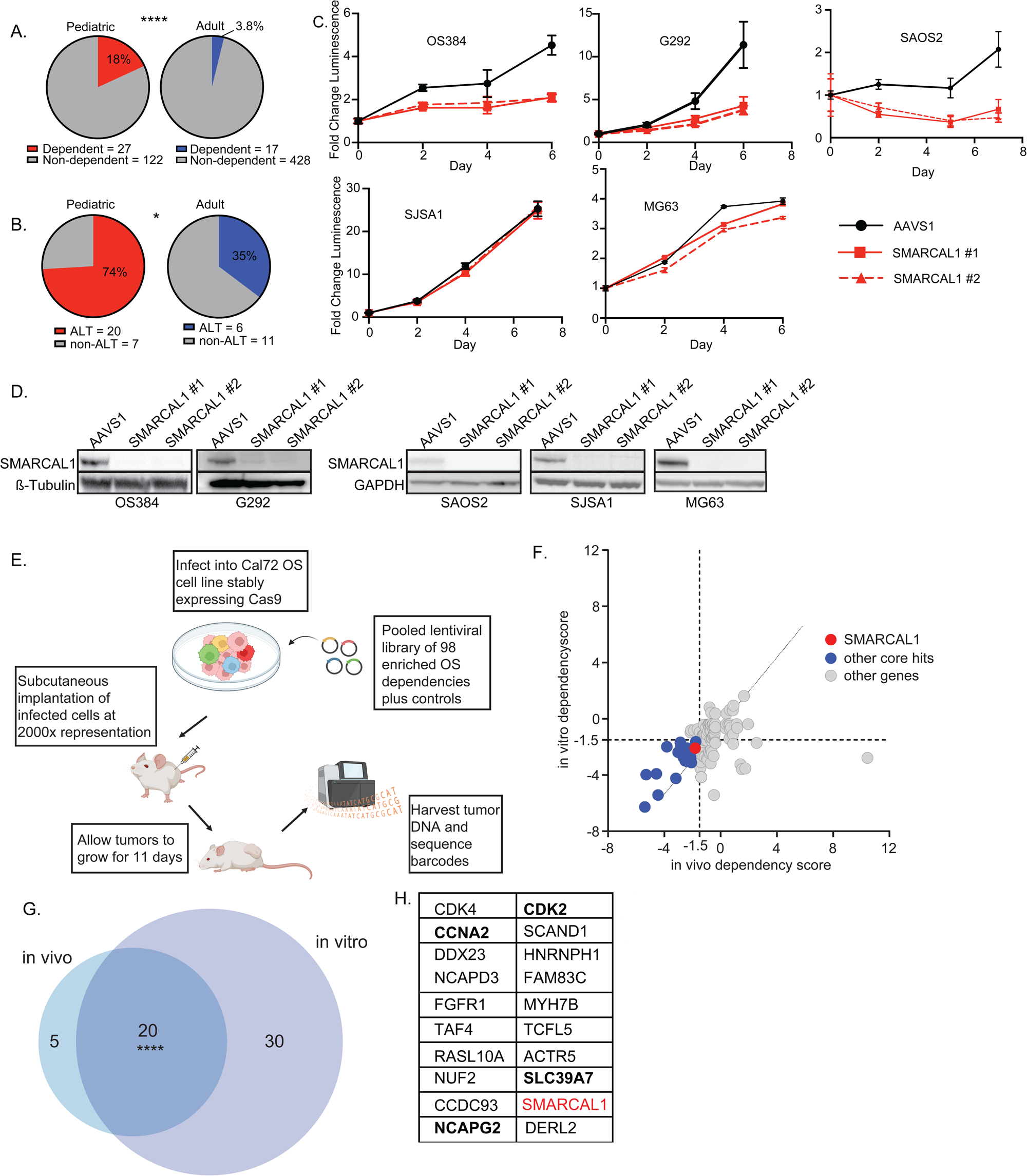
*SMARCAL1* is a top target in ALT+ osteosarcoma *in vitro* and scores in a pooled *in vivo* CRISPR screen. **a.** Pie charts demonstrating proportion of *SMARCAL1-*dependent cell lines in pediatric versus adult cancer models in DepMap. Total numbers of percent breakdowns below. Pediatric cancers show a significantly higher rate of *SMARCAL1* dependency (18%; 27/149) compared to adult cancers (3.8%; 17/454). (****p < 0.0001 by Fisher’s exact test. **b.** Pie charts demonstrating ALT status distribution (by ALTitude prediction) among *SMARCAL1*-dependent cell lines stratified by age group. Among *SMARCAL1*-dependent lines, pediatric models are significantly enriched for ALT-positive status (74%; 20/27) compared to adult models (35%; 6/17). *p < 0.05 by Fisher’s exact test. **c.** Viability assay demonstrating *SMARCAL1* CRISPR KO (*SMARCAL1* #1, #2) vs. non-targeting (AAVS1) in ALT+ osteosarcoma cell lines OS384, G292 and SAOS2 and of non-ALT cell lines SJSA1 and MG63. X-axis represents time point of luminescence measurement, y-axis represents the fold change in luminescence **d.** Western immunoblotting confirming *SMARCAL1* CRISPR KO in the osteosarcoma cell lines shown in **c** 3-5 days post puromycin selection. Two guides targeting *SMARCAL1*(#1, #2) and the control (AAVS1) are shown. GAPDH and β-Tubulin are used as loading controls. **e.** Schematic showing *in vivo* screen performed in the Cal72 ALT+ osteosarcoma cell line. A pooled library of 98 enriched osteosarcoma (OS) guides was used, plus controls (positive and negative) in stably Cas9-expressing cells. Figure made using Biorender. **f.** Scatter dotplot depicting the dependency scores identified by STARS (see methods) for the genes screened: day 11 *in vivo* (x-axis) vs. day 25 *in vitro* (y-axis). Dots represent the 112 out of the 118 genes in the pooled screen that received a STARS dependency score. Significance cutoffs: STARS score ≤ -1.5, p-value ≤ 0.10. The common gene hit *SMARCAL1* is highlighted red. The other common gene hits are highlighted blue. **g.** Venn diagram presenting the overlap between the dependency gene hits identified by STARS for day 11 *in vivo* vs. day 25 *in vitro* in the pooled screen. Overlap significance **** p < 0.0001 by two-tailed Fisher exact test. **h.** list of genes scoring as common gene hits. *SMARCAL1*is in red. Common essential genes are bolded in black.

We next tested whether *SMARCAL1* dependency persisted *in vivo* and whether it outperformed other candidate osteosarcoma liabilities using a pooled CRISPR screening approach. We first identified 98 gene dependencies enriched in osteosarcoma models compared to all other DepMap cell lines, agnostic to ALT status. We designed a single-guide RNA (sgRNA) lentiviral CRISPR KO library with four guides per gene and included intronic guides to control for copy number-related toxicity effects, pan-essential gene guides as positive controls, and non-targeting and intergenic guides as negative controls (**Extended Data Table 7**). We performed a LOF screen using this library in a subcutaneous xenograft model of the ALT+ osteosarcoma cell line Cal72,^40^ with a parallel screen performed *in vitro* to compare performance of our custom library relative to the DepMap genome-scale screen. The correlation between the *in vitro* and *in vivo* screens was high (Pearson *r* = 0.60, *P* < 1.0 × 10⁻⁴), and *SMARCAL1* depletion led to a significant viability effect *in vivo* (STARS algorithm score = -1.807, *P* = 0.05, see **Methods**). We identified 20 gene dependency hits that scored at both *in vitro* day 25, a time point similar to the DepMap screen timing, and *in vivo* day 11, a timepoint chosen to avoid clonal expansion of negative controls *in vivo* (**Fig. 2e-g**). As expected, several pan-essential genes, included in the screen as positive controls, also scored strongly (bolded in **Fig. 2h**). Taken together, these results validated *SMARCAL1* as a reproducible ALT-associated dependency and prioritized it relative to other osteosarcoma-enriched dependencies in an ALT+ *in vivo* model.

### ALT status predicts *SMARCAL1* dependency in pediatric neuroblastoma

Given the high prevalence of ALT in pediatric tumors, we next asked whether ALT status predicted *SMARCAL1* dependency in neuroblastoma, another common and devastating pediatric cancer. In older children with high-risk neuroblastoma, tumors often display ALT along with *ATRX* alterations, including in-frame fusions (IFFs) (explored in detail in the next section). These children have a long-term survival of <50% and no available targeted treatments.^12,41,42^ Furthermore, ALT is prevalent in relapsed neuroblastoma, which has a particularly poor prognosis. We examined differential dependencies in the three ALT+ neuroblastoma cell lines profiled with DepMap CRISPR screening, two confirmed to be ALT+ with IFFs in *ATRX* (SKNMM and CHLA90) that are previously well-described^43,44^ and one ALT+ line with WT *ATRX* (SKNFI). (**Extended Data Table 1, Extended Data Fig. 7a-b**). We examined the enriched dependencies in these lines compared to all other DepMap cell lines, and strikingly, we saw that *SMARCAL1* scored as one of the top enriched dependencies (**Fig. 3a**). We used CRISPR to KO *SMARCAL1* in the *ATRX* IFF ALT+ cell lines CHLA90 and SKNMM, and the *ATRX* WT cell line SKNFI and used CTG to measure the effect on cell viability, surprisingly observing a growth effect only in the *ATRX* altered cell lines (**Fig. 3b-c**). We sought an explanation for this and profiled protein expression levels of SMARCAL1 in SKNFI and several outgroup cell lines. We observed that while the transcript level of *SMARCAL1* is comparable by RNA-seq to other DepMap cell lines, it expresses very low SMARCAL1 protein (**Extended Data Fig. 7c**). Thus, we hypothesize that this low protein level may cause ALT in SKNFI, similar to other ALT+ tumors with *SMARCAL1* loss,^45–47^ explaining why these cell lines are resistant to *SMARCAL1* KO, despite being ALT+.

**Fig. 3.**
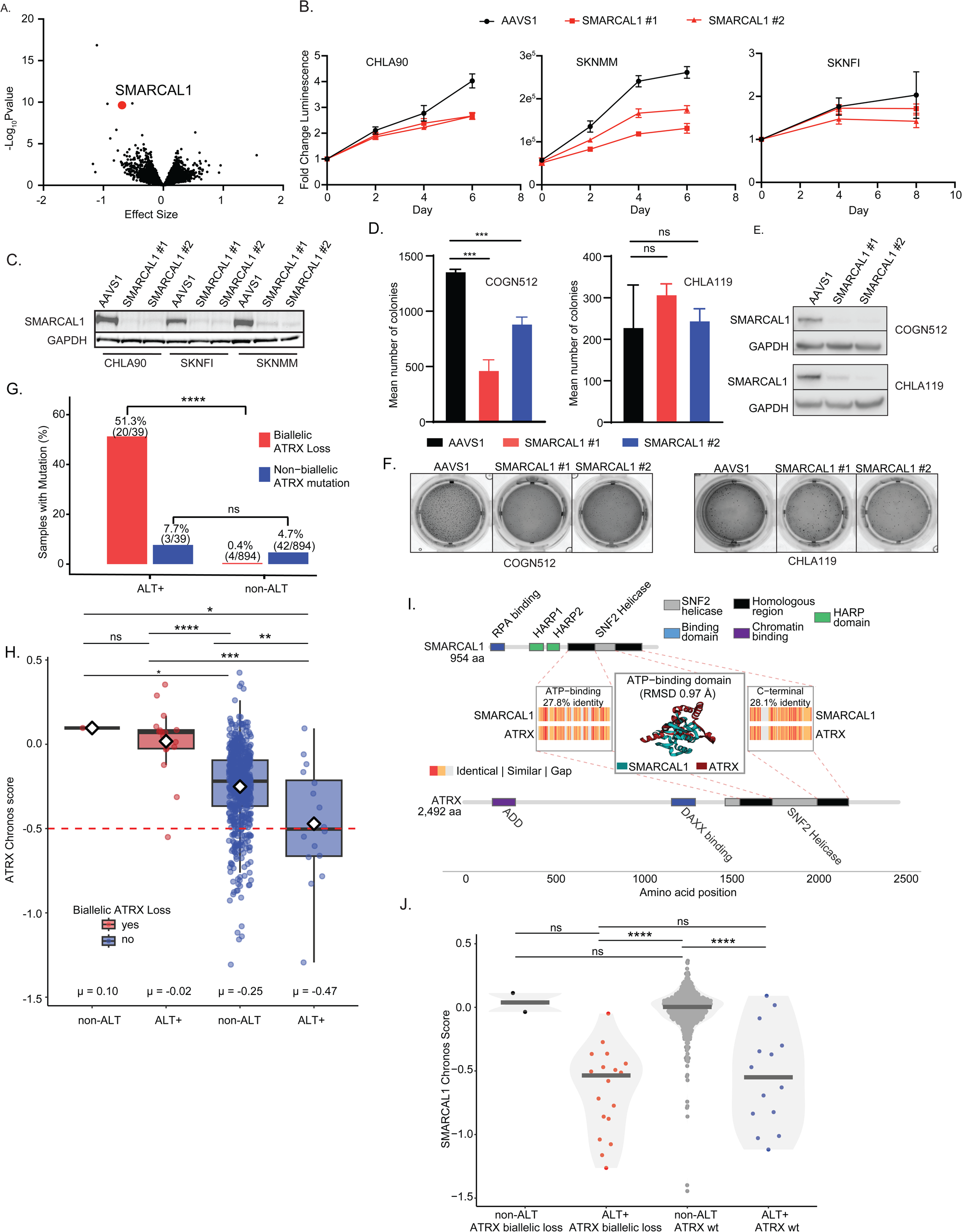
*SMARCAL1* is a dependency in neuroblastoma, and integration of *ATRX* mutation calls with ALT predictions suggests differential mechanisms of ALT-specific dependencies. **a.** Volcano plot showing a two-class comparison analysis between ALT-positive neuroblastoma models (3) and all other cell lines in DepMap (1183). Effect size (x-axis) is plotted against statistical significance (-log_10_ p-value, y-axis). *SMARCAL1* (red) is highlighted. **b.** Viability assay demonstrating *SMARCAL1* CRISPR KO (*SMARCAL1* #1, #2) vs. non-targeting (AAVS1) in ALT+ neuroblastoma cell lines CHLA90, SKNMM and SKNFI. X-axis represents time point of luminescence measurement; y-axis represents the fold change in luminescence. **c.** Western immunoblotting confirming *SMARCAL1* CRISPR KO in the neuroblastoma cell lines shown in **b**. 3-5 days post puromycin selection. Two guides targeting *SMARCAL1*(#1, #2) and the control (AAVS1) are shown. GAPDH is used as a loading control. **d.** Bar graphs depicting number of colonies in a methylcellulose anchorage-independent growth assay following CRISPR KO of *SMARCAL1* in COG512 ALT+ and in CHLA119 non-ALT cell lines. Two guides*SMARCAL1*(#1, #2) and the control (AAVS1) are shown. Results were performed in triplicate for COGN512 and in duplicate for CHLA119. ***p<0.0005 by unpaired T-test. **e.** Western immunoblotting confirming *SMARCAL1* KO in the neuroblastoma cell lines shown in **d**. GAPDH is used as a loading control.**f.** Representative images of colony formation assay shown in **d-e**. **g.** Percentage of cell lines with biallelic *ATRX* loss in ALT+ (n=39) versus non-ALT (n=894) groups as classified by ALTitude. ****p<0.0001 by Fisher’s exact test. **h.** Bar and whisker plot demonstrating *ATRX* Chronos score (y-axis) grouped by non-ALT vs. ALT+ by ALTitude predictions and biallelic loss of *ATRX* (red) vs. WT (blue). Pairwise comparisons were performed using two-sided Wilcoxon rank-sum tests with Benjamini-Hochberg correction (ns, p ≥ 0.05; *p < 0.05; **p < 0.01; ***p < 0.001; ****p < 0.0001). Diamond indicates group mean; μ values shown below each group. **i.** Schematic showing domain architecture of SMARCAL1 and ATRX proteins. SMARCAL1 contains RPA-binding, HARP1, HARP2, and helicase domains. ATRX contains ADD, DAXX-binding, helicase, and HP1-binding domains. Scale indicates amino acid position. j. *SMARCAL1* Chronos scores stratified by ALT status and *ATRX* bi-allelic mutation status. More negative scores indicate stronger dependency. ALT-positive cell lines show significantly greater *SMARCAL1* dependency regardless of *ATRX* status ****p<0.0001 by Wilcoxon rank-sum tests.

We then asked if we could infer *SMARCAL1* dependency based on ALT+ status in cancer models outside of the DepMap. To do this, we used two patient-derived low-passage neuroblastoma models, specifically, COGN512 and CHLA119. COGN512 is confirmed as ALT+ by C-circle analysis,^44^ and CHLA119 is non-ALT.^12^ We confirmed both by telomere FISH (**Extended Data Fig. 7d**). As these cells are low-passage and contain both adherent and suspension cells in culture, methylcellulose assays were used to probe for viability effects of *SMARCAL1* CRISPR KO. We found that *SMARCAL1* depletion led to significantly reduced anchorage-independent growth in the ALT+ line COGN512. Conversely, we saw no growth effects of *SMARCAL1* loss in the non-ALT cell line CHLA119 (**Fig. 3d-f**). These data show that ALT status can predict *SMARCAL1* dependency and support *SMARCAL1* as a rational therapeutic target candidate for multiple ALT+ cancers.

### Reanalysis of DepMap WGS data identifies numerous new *ATRX* loss-of-function cell line models, driven by diverse, complex and cryptic mutational events

How cancers establish and sustain ALT is not fully understood, though LOF mutations in the chromatin remodeler *ATRX*, or more rarely, its binding partner, *DAXX*, are thought to drive a substantial subset of ALT+ malignancies.^48,49^ However, the true rates of *ATRX* loss in cancer cell lines, including across DepMap, remain unknown. This is due to the complexity and diversity of the genomic alterations reported to target *ATRX*, including structural variants (SVs) and complex insertions/deletions (Indels), which are notoriously difficult to resolve by short read DNA-sequencing alone.^41,49,50^ We thus re-analyzed the DepMap WGS data using clinical-genomics-grade pipelines, calling single/multiple nucleotide variants (SNV/MNVs), copy number variants (CNV), SVs and Indels (see **Methods**). Our complete set of ATRX loss calls, and underlying justifications for each call, are presented in **Extended Data Table 1**.

Excluding synonymous mutations, we found 138 DepMap cell lines carry at least one candidate *ATRX*-inactivating variant, but most do not correspond to complete loss of canonical ATRX function (e.g. lacking clonal stop-gained, frameshift, deletions; evidence of inactivation of both alleles if female). Indeed, we only identified a total of 24 cell lines exhibiting likely complete canonical LOF, of which 16 have not been previously reported in the literature (known cell lines were osteosarcoma *n* = 5, neuroblastoma *n* =2 and AML *n* = 1 **Extended Data Table 8**). A subset of these mutations (e.g. frameshifts) would be expected to lead to loss of detectable ATRX protein, which we confirmed by Western blot for 11 cell lines, (**Extended Data Table 1**, **Extended Data Fig. 7e-g;** 100% concordance with our curated *ATRX* loss assignments based on genomics).

Our *ATRX* loss cell lines highlighted striking mutational/mechanistic diversity, with many events difficult to capture without manual review. For example, in addition to numerous canonical protein-truncating mutations (e.g. frameshift variants), we observed: (i) out-of-frame multi-exon deletions (e.g. NP8); (ii) near whole-gene deletions (e.g. the osteosarcoma line OS384, small cell lung cancer line NCIH1184, and myxofibrosarcoma line NRHMFS2); (iii) promoter or 5′ regulatory deletion abrogating transcription (e.g. NRHUPS4, an undifferentiated pleomorphic sarcoma model); (iv) complex rearrangements including in-frame structural events removing most of the gene (e.g. neuroblastoma lines SKNMM and CHLA90 – previously reported^41,43^); and (v) point mutations inactivating the ATRX helicase ATPase domain (e.g. DMS153, a small cell lung cancer line, and OS525, an osteosarcoma model). Taken together, these results underscore that “mutated *ATRX*” is not synonymous with “complete loss of *ATRX*,” and that defining *ATRX* loss based on genomics requires intense manual curation and integration of allele dosage, sex/X copy state, and functional readouts like *ATRX* dependency status and gene expression. Notably, our final calls include several, as far as we know, first-reported *ATRX* loss disease models for myxofibrosarcoma (NRHMFS2), undifferentiated pleomorphic sarcoma (NRHUPS2, NRHUPS4), small cell lung cancer (DMS153, NCIH1105) and rhabdomyosarcoma (RHJT, RH4), providing a resource for studying *ATRX* loss in these diverse contexts.

### Integration of *ATRX* loss, ALT calls, and dependency profiles in DepMap provides a potential mechanistic basis for ALT-specific dependencies

We next used these new mutation calls to re-visit the relationship between ALT and *ATRX* loss, at unprecedented scale. Consistent with expectation, 20/24 (83%) of our *ATRX* loss cell lines were also called ALT+ by ALTitude (**Fig. 3g),** supporting the validity of our mutation calls via an independent functional readout, and consistent with the idea that complete loss of canonical ATRX function is required to contribute to ALT. The existence of four *ATRX* loss cell lines, which do not exhibit ALT (all adult males, three with small cell lung cancer and one with AML), suggests that these are either passenger events, or that *ATRX* loss can drive cancer via ALT-independent mechanisms. Of the remaining 19 *ATRX*-intact ALT+ cell lines, two models, G292 & OS052, carry likely pathogenic *DAXX* loss variants and two others, the aforementioned SKNFI (at the protein level, see **Extended Data Fig. 7c**), and NY (biallelic deletion, previously reported), have loss of SMARCAL1 (**Extended Data Table 1**).^40^ However, the remaining 15 ALT+ cell lines were *ATRX/DAXX/SMARCAL1* WT, suggesting one or more unknown drivers of ALT (see **Discussion**). Interestingly, we also found that *ATRX* itself was a strong and selective dependency in ALT+ cell lines, when ALT was *not* driven by *ATRX* loss (**Fig. 3h**). Conceptually, this is consistent with functional compensation, for example, such that loss of one chromatin-associated factor (e.g. ATRX), which drives ALT initially, shifts reliance onto others (e.g., SMARCAL1), and vice versa. Notably, both *ATRX* and *SMARCAL1* encode Sucrose Non-Fermenting (SNF) 2 family members and ATP-dependent DNA chromatin remodelers, and we found that these exhibit significant structural and sequence conservation in the helicase domain (**Fig. 3i**, see **Methods**).^51^ We also observed a single ALT+ cell line (OS525) with a likely inactivating mutation targeting this specific domain of ATRX (**Extended Data Table 1**). For both ATRX and SMARCAL1, these domains act at stalled replication structures, which often arise at regions characterized by repetitive DNAsequences, such as telomeres.^52–55^ Thus, this shared function potentially underpins a classical synthetic lethal relationship and provides a plausible mechanistic basis for *SMARCAL1* dependency in a sub-set of ALT.

We next revisited the relationship between *ATRX* status, ALT, and dependence on *SMARCAL1*, hypothesizing that *ATRX* loss may potentiate *SMARCAL1* dependency in ALT. Surprisingly however, *SMARCAL1* dependency did not differ detectably between *ATRX* loss (*n = 19*) and *ATRX* WT (*n* = 14) ALT+ groups (**Fig. 3j**; Cohen’s d = -0.3001, *Padj* = 0.5). Thus, *SMARCAL1* dependency appears to be a general feature of ALT, regardless of whether ALT is driven by *ATRX* loss or some other event(s). We observed, however, that *ATRX/DAXX* loss, without taking into account ALT status, was markedly enriched among *SMARCAL1*-dependent lines relative to non-dependent lines (37.8% vs. 0.64%; Fisher’s exact test, *P* =2.3 × 10^-18^); a total of only four cell lines in our DepMap dataset were *ATRX* lost or *DAXX* mutant but not *SMARCAL1* dependent (**Extended Data Fig. 8**). We note that recently, *SMARCAL1* was observed as a dependency in ALT+ high-grade gliomas specifically in the absence of *ATRX*.^33^ In the DepMap dataset, we had only two high-grade gliomas which were ALT+ by ALTitude; these were both *ATRX* lost and *SMARCAL1* dependent, thus limiting our ability to examine dependency in this disease context outside of *ATRX* loss (**Extended Data Table 1**). When examining specific lineages in our dataset enriched for *SMARCAL1* dependency, we observed the clearest *ATRX* genotype to *SMARCAL1* dependency relationship in osteosarcoma models. *ATRX* loss is a common finding in osteosarcomas due to deletions or complex rearrangements, with an estimated prevalence of 20-40%, and is associated with ALT+.^56–60^ Therefore, we hypothesized that SMARCAL1 is playing a distinct mechanistic role in this disease, similar to high-grade gliomas, with the absence of *ATRX/DAXX*.

### ATRX and SMARCAL1 compensate functionally in ALT+ cancer cells

We next explored these ideas mechanistically, again in the context of osteosarcoma models. We first asked whether SMARCAL1 expression modified the consequence of *ATRX* loss in a cell line that lacked endogenous *SMARCAL1*. We used the NY osteosarcoma line, which carries a biallelic deletion of *SMARCAL1*, is ALT+, and retains wild-type *ATRX*.^40^ We constitutively expressed SMARCAL1 in NY cells using a lentiviral construct and compared growth with a GFP overexpression control. SMARCAL1 overexpression reduced proliferation by IncuCyte® live cell imaging, which suggested that, at least in this model, simultaneous presence of functional SMARCAL1 and ATRX imposed a fitness cost in this ALT+ context (**Fig. 4a-b**). We next disrupted ATRX by CRISPR in both SMARCAL1-expressing and GFP-expressing NY cells. In GFP controls, *ATRX* KO also reduced fitness, whereas SMARCAL1 expression mitigated this effect and partially restored proliferation (**Fig. 4a-c**). These results indicated that SMARCAL1 buffered the consequences of ATRX loss.

**Fig. 4.**
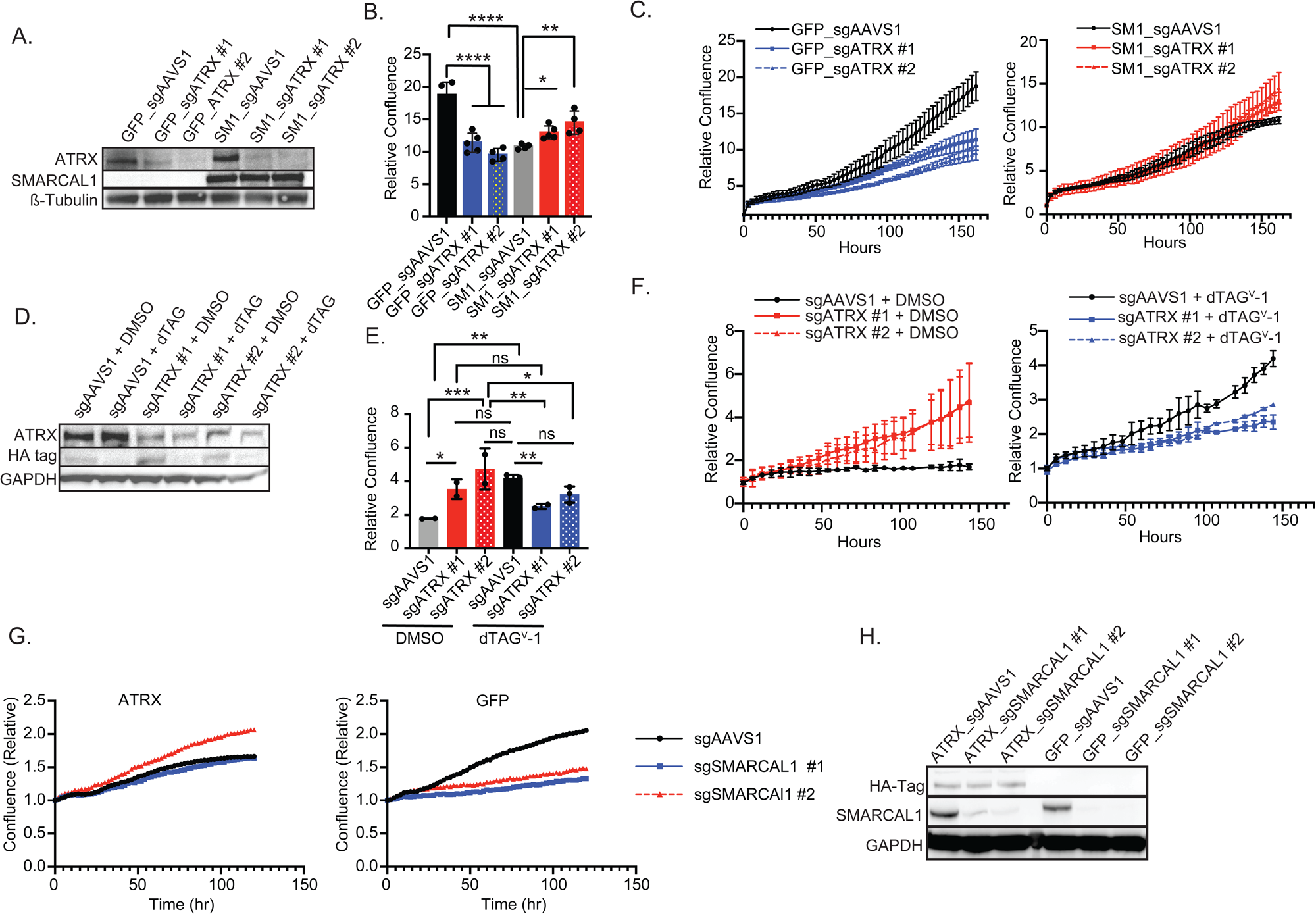
SMARCAL1 and ATRX functionally compensate for optimal viability in ALT+ osteosarcoma and neuroblastoma. **a.** Western immunoblotting showing SMARCAL1 and ATRX protein levels in the osteosarcoma cell line NY. *SMARCAL1* and a control (GFP) overexpression and CRISPR KO of *ATRX* with two guides (#1, #2) and a control (AAVS1) are shown. β-Tubulin is used as a loading control. **b.** Bar graphs showing statistical analysis of final time point of IncuCyte® life cell imaging shown in **c** in NY osteosarcoma cells that overexpress *SMARCAL1* (grey/red) or GFP (black/blue) and have CRISPR KO of *ATRX* (blue and red). * p < 0.05, ** p<0.005, **** p<0.0001 by Brown-Forsythe and Welch ANOVA. **c.** Live cell imaging counts depicting relative confluence of NY cells that overexpress *SMARCAL1* or a GFP control and concurrently have KO of *ATRX* (#1, #2) or gene desert control (AAVS1). X axis shows time in hours; y axis represents the relative confluence normalized to time zero. **d**. Western immunoblotting demonstrating *ATRX* KO and/or SMARCAL1 protein degradation (HA tag antibody) after dTAG^v^-1 treatment of NY cells for 24 hours. GAPDH is used as a loading control. **e.** Live cell imaging counts depicting relative confluence of NY osteosarcoma cells infected with FKBP12^F36V^-SMARCAL1 treated with 1µM dTAG^V^-1. X-axis depicts hours post-treatment; y-axis depicts confluence normalized to time zero. **g.** Graphs showing viability of OS384 cells with ATRX vs. GFP overexpression infected with sgRNAs against SMARCAL1 (#1 & #2) or AAVS1 by live cell imaging over 120 hours. X-axis represents time in hours; y-axis represents relative confluence to time zero. **h.** Western immunoblotting showing levels of ATRX overexpression (HA-tag antibody) vs. overexpression control with CRISPR KO of *SMARCAL1* with two guides (#1, #2) or a control (AAVS1) in OS384 osteosarcoma cells. GAPDH is used as a loading control.

To confirm whether this rescue depended directly on SMARCAL1 protein, we used our FKBP12^F36V^ degron system to acutely degrade exogenous SMARCAL1 protein. We expressed FKBP12^F36V^-tagged SMARCAL1 in NY cells, treated cells with dTAG^V^-1, and monitored growth by live cell imaging. Acute SMARCAL1 degradation partially reversed the rescue phenotype, which supported a direct, on-target relationship between SMARCAL1 protein abundance and the ability to tolerate *ATRX* loss (**Fig. 4d-f**).

We then performed the reciprocal experiment in an ALT+ background that lacked *ATRX*. We used the OS384 osteosarcoma line, which carries a complete *ATRX* deletion and expresses neither ATRX transcript nor protein (**Extended Data Fig. 7f**). We introduced full-length wild-type *ATRX* by lentiviral transduction and performed *SMARCAL1* CRISPR KO. In OS384 cells with *ATRX* re-expression, the dependency on *SMARCAL1* was rescued, with equal or superior growth seen with SMARCAL1 KO in the presence of *ATRX* overexpression by IncuCyte® live cell imaging (**Fig. 4g-h**). Together, these data support a model in which ALT+ cells tolerate disruption of one of these remodelers when the other remained functionally available, and they further suggest that the combined loss or presence of both proteins may create a fitness bottleneck in ALT+ cells.

### *SMARCAL1* KO leads to overexpression of ALT-associated DNA damage-related genes in an ALT+ cell line, but not in non-ALT cells

We were interested in further unraveling the mechanistic basis for *SMARCAL1*’s selective dependency in ALT. We reasoned that gene expression changes occurring upon *SMARCAL1* KO, and their relationship to ALT, could provide a roadmap towards the specific cellular processes underpinning the selective fitness effect. We first integrated our ALT calls with RNA-seq data from 376 DepMap cell lines from ALT-relevant lineages (39 ALT+ and 337 ALT−), comparing genome-wide gene expression in ALT+ vs. all other cell lines. Notably, at the pathway level, gene set enrichment analysis (GSEA) revealed similar enrichment scores for our assay-confirmed ALT+ cell lines vs the new ALTitude-predicted ALT calls (**Fig. 5a-b**), supporting the ALTitude calls in an independent molecular modality. From the differential expression results, we defined a set of differentially expressed telomere maintenance genes by intersecting significant differentially expressed genes with curated telomere-related gene sets from MSigDB (**Extended Data Table 9**). This yielded 87 genes upregulated in ALT+ cells (ALT_up) and 28 genes downregulated in ALT+ cells (ALT_down). These 2 resulting gene sets included well-established ALT-associated factors, known to also play a role in replicative stress and DNA damage response, including *BLM*, *BRCA2*, *RAD51*, *XRCC3^6134,62-65^*⍰ in ALT_up, as well as genes expected to be depleted in ALT contexts such as *ATRX* and *TERT*, as well as *TP53* and *RB1* in ALT_down.

**Fig. 5.**
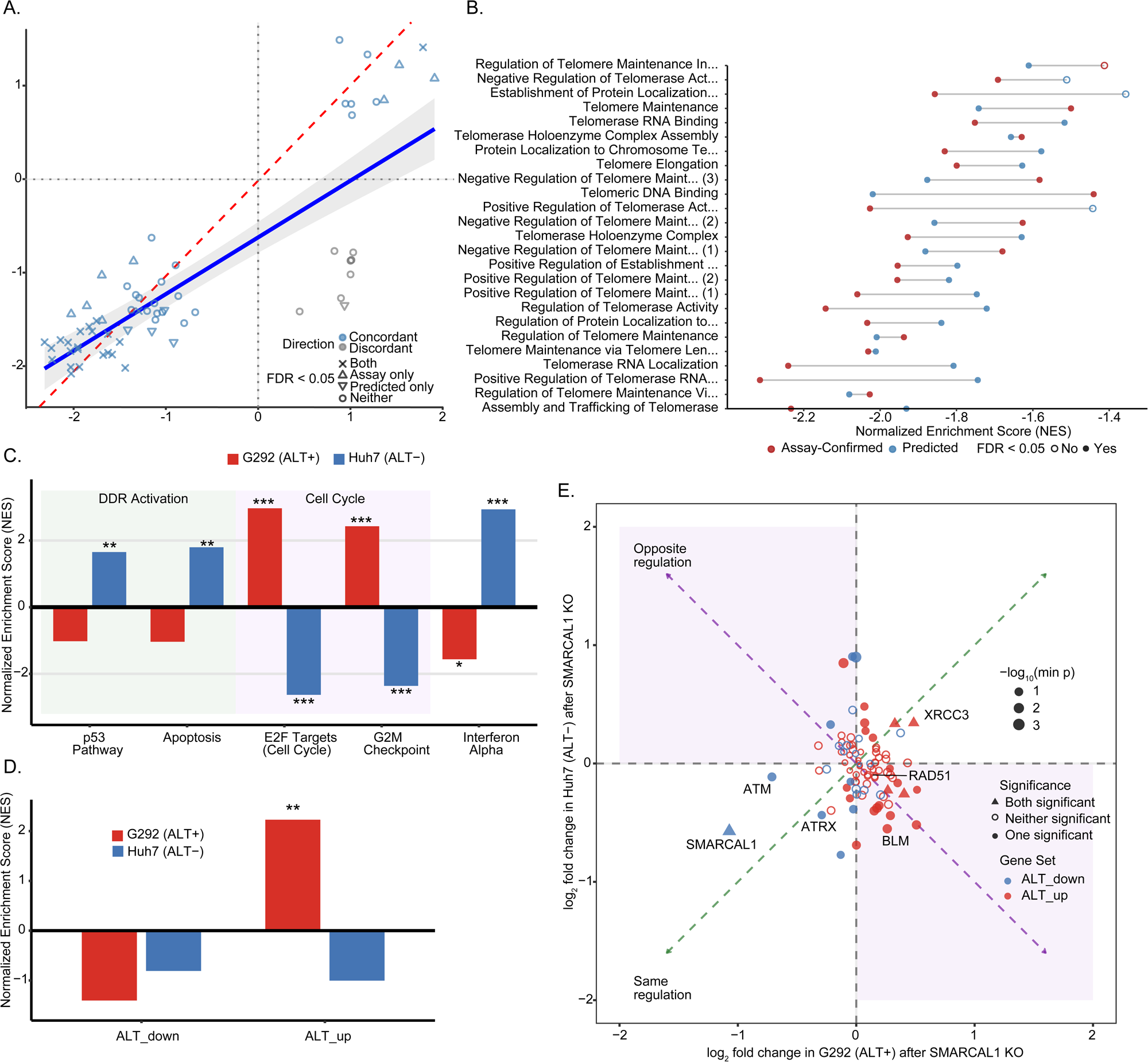
Validation of ALT pathway signatures and differential response to *SMARCAL1* knockout in ALT-positive versus ALT-negative cell lines. **a.** Concordance of pathway enrichment between assay-confirmed and ALTitude-predicted ALT-positive samples. Each point represents a gene set, with normalized enrichment scores (NES) from assay-confirmed ALT+ samples (x-axis) plotted against NES from predicted ALT+ samples (y-axis). Blue points indicate concordant directionality; grey points indicate discordant results. Pearson correlation (r = 0.8) and proportion of gene sets with matching direction (56/66, 85%) are shown. Solid blue line represents linear regression fit with 95% confidence interval (shaded region); dashed red line indicates y = x. **b.** Comparison of Normalized Enrichment Scores (NES) for telomere and telomerase-related gene sets between assay-confirmed (red) and predicted (blue) ALT-positive classifications. Negative NES values indicate depletion of these gene sets in ALT-positive relative to non-ALT samples. **c.** Hallmark pathway enrichment following *SMARCAL1* KO in G292 (ALT+, red) versus Huh7 (non-ALT, blue) cell lines. Pathways are grouped by DNA damage response (DDR) activation and cell cycle regulation. Significance: *p < 0.05, **p < 0.01, ***p < 0.001 by Benjamini-Hochberg correction. **d.** NES for ALT-associated gene sets (ALT_down and ALT_up) following *SMARCAL1* KO in G292 and Huh7 cells. **p < 0.01. **e.** Comparison of ALT gene regulation following *SMARCAL1* knockout between G292 (ALT+, x-axis) and Huh7 (ALT−, y-axis) cell lines. Each point represents a gene from ALT_down (blue) or ALT_up (red) gene sets. Point shape indicates significance status: triangles denote genes significant in both cell lines, filled circles indicate significance in one cell line only, and open circles represent genes not significant in either line. Point size is scaled by −log_10_(minimum p-value across both comparisons). Quadrants are annotated to indicate concordant regulation (green dashed diagonal; genes changing in the same direction in both lines) versus opposite regulation (purple shading; genes with discordant responses). A lack of correlation (r = −0.014, p = 0.92) is shown, indicating that ALT-associated genes respond differently to *SMARCAL1* loss depending on ALT status, with ALT_up genes (red) showing preferential upregulation specifically in the ALT+ context (Q4, lower right quadrant). Key genes involved in telomere maintenance and DNA repair are labeled.

*SMARCAL1* has been reported to influence transcriptional programs in mammalian cells, but this has not previously been explored in the ALT context.^62–64^ To determine how *SMARCAL1* loss alters gene expression specifically in the setting of ALT, we next performed RNA-seq following CRISPR KO of *SMARCAL1* in the G292 osteosarcoma cell line. G292 is an ALT+ model with a *DAXX* rearrangement^40,65^ and shows strong *SMARCAL1* dependency both in DepMap and in our validation experiments (see **Fig. 2c-d**, **Extended Data Fig. 6c-f**). As a lineage-mismatched and non-ALT comparator with minimal *SMARCAL1* dependence, we also analyzed RNA-seq data from Huh7 hepatocellular carcinoma cells following CRISPR KO of *SMARCAL1* (GSE221372).^66^ Huh7 carries a *TERT* promoter mutation and is telomerase-positive.^67^ We used GSEA (see Methods) to summarize the transcriptional consequences of SMARCAL1 loss, focusing on MSigDB Hallmark gene sets and our DepMap-derived ALT_up/ALT_down signatures. Strikingly, SMARCAL1 KO produced diametrically opposed transcriptional responses in ALT+ versus non-ALT cells across proliferation and immune-signaling programs: E2F targets (NES = +2.97 vs −2.63) and G2M checkpoint (NES = +2.43 vs −2.36) were upregulated in G292 but downregulated in Huh7, whereas interferon-α and interferon-γ responses showed the opposite pattern (IFN-α NES = −1.56 vs +2.94; IFN-γ NES = −1.38 vs +2.49). Non-ALT Huh7 cells further activated canonical stress-response programs, including the P53 pathway, apoptosis, and TNFα/NFκB signaling, whereas ALT+ G292 cells failed to induce these pathways (Fig. 5C). Thus, although SMARCAL1 loss is known to induce replication stress across all cells, the downstream transcriptional response diverges sharply in ALT.

We next asked whether SMARCAL1 perturbation modulates the ALT-associated transcriptional program itself. In G292 cells, our DepMap-derived ALT_up genes were strongly enriched following *SMARCAL1* KO (NES = +2.24, P = 5.7 × 10⁻⁷), consistent with amplification of the ALT-associated transcriptional signature (**Fig. 5d**, **Extended Data Table 10-11**). In contrast, ALT_up was not enriched in the non-ALT Huh7 cells following *SMARCAL1* KO (NES = −1.00, P = 0.46). Among the genes driving the ALT-specific response, *BLM*, *RAD54L*, and *NCL* were all significantly downregulated in Huh7 (logFC = −0.55, −0.52, and −0.57, respectively; all P < 0.05) but trended upward in G292 (logFC = +0.26, +0.51, and +0.18) (Fig. 5e).^68^ These data suggest that SMARCAL1 KO does not have an entirely benign effect in non-ALT cells, but rather that its loss can be tolerated due to the function of compensatory cellular machinery. However, in ALT+ cells, SMARCAL1 appears to play a crucial role in balancing key ALT effector genes; SMARCAL1 loss in ALT+ cells leads to further upregulation of the ALT program and subsequent cell death, perhaps due to lack of other compensatory cell mechanisms, such as, in some circumstances, ATRX or DAXX.

### *SMARCAL1* loss significantly augments C-circles and leads to G2/M checkpoint activation and aneuploidy in ALT+ models

To experimentally investigate the molecular underpinnings of *SMARCAL1* dependency in ALT, suggested by our RNA-seq analysis, we next queried the effects of SMARCAL1 depletion on features of ALT and replication stress as well as cell cycle and DNA damage *in vitro*. We first asked how SMARCAL1 depletion affects C-circles in three osteosarcoma and two neuroblastoma cell lines, using both CRISPR KO and degron systems. With SMARCAL1 depletion, C-circles have previously been reported to increase even in telomerase positive models.^31^ We found that only *SMARCAL1* KO or degradation in ALT+ cell lines that were dependent on *SMARCAL1* led to significantly increased C-circles. There was no significant change in the SMARCAL1-low ALT+, neuroblastoma line SKNFI, nor in the non-ALT model MG63 (**Fig. 6a**). We then tested whether the functional compensation between SMARCAL1 and ATRX also extended to ALT molecular readouts. We used the NY osteosarcoma rescue system and measured C-circles after manipulating SMARCAL1 and ATRX. When we overexpressed *SMARCAL1* in NY cells, we reduced C-circles markedly, consistent with prior reports that *SMARCAL1* reconstitution can suppress ALT features in *SMARCAL1*-null models.^45^ Interestingly, when we performed KO of *ATRX* in the *SMARCAL1*-overexpressing background, we partially restored C-circle positivity (**Fig. 6b**). These data support a model in which a subset of ALT+ cancers maintain an optimal ALT state when cells retain either functional SMARCAL1 or ATRX, and that this state is lost when these levels are perturbed.

**Fig. 6.**
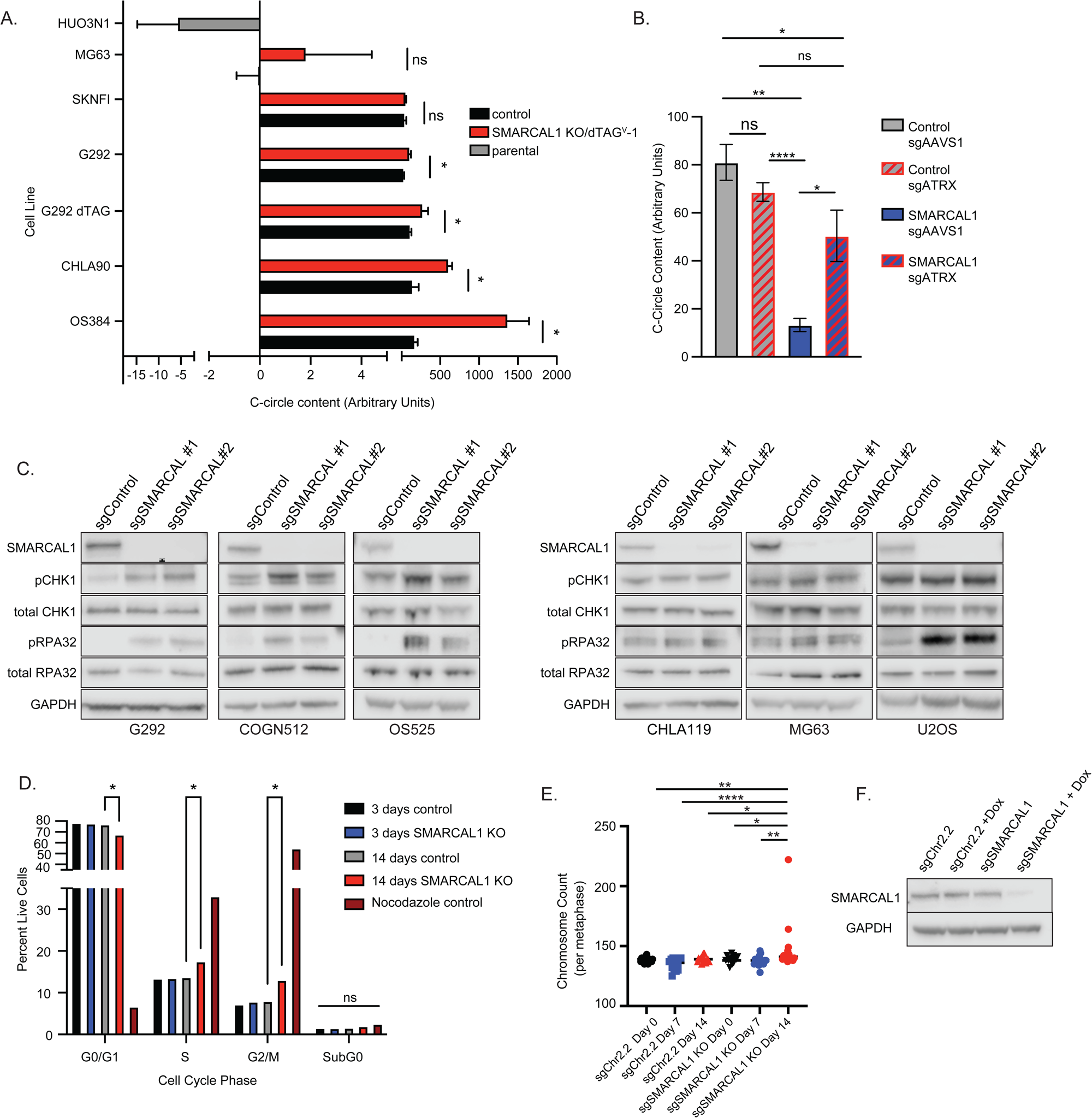
*SMARCAL1* KO in ALT+ cells leads to increased c-circle formation, G2/M phase arrest and aneuploidy. **a.** Bar graphs demonstrating calculated C-circle content by qPCR performed after perturbation (CRISPR KO or 1µM dTAG^V^-1 for 24 hours) in neuroblastoma (SKNFI, CHLA90, SKNFI) and osteosarcoma (HUO3N1, MG63, G292, OS384) cell lines. C-circle content greater than 5 is considered ALT+. HUO3N1 parental cell line is included to demonstrate a parental non-ALT cell line control. * p<0.05 by unpaired T-test with Welch’s correction. **b.** Bar graphs demonstrating calculated C-circle content by qPCR in NY cells after *SMARCAL1* vs. control overexpression and/or and *ATRX* CRISPR KO. *SMARCAL1* overexpression significantly reduces the C-circle content compared to controls, and *ATRX* KO partially reverses this effect. *<0.05, ** p<0.005, **** p<0.0001 by unpaired T-test with Welch’s correction. **c.** Western immunoblotting depicting *SMARCAL1-*dependent osteosarcoma (G292, OS525) and neuroblastoma (COGN512) cell lines as well as *SMARCAL1* non-dependent osteosarcoma (MG63, U2OS) and neuroblastoma (CHLA119) cell lines demonstrating the effect of perturbation with CRISPR KO of *SMARCAL1* (#1, 2) vs control KO on total and pCHK1 and total and pRPA32. GAPDH is used as a loading control. **d.** Cell cycle analysis of OS384 cells infected with a doxycycline-inducible CRISPRi system targeting *SMARCAL1* vs. a gene desert control (Chr.2.2). The graph represents the percent live cells in the different cell cycle phases after 3 and 14 days following doxycycline induction. Cells treated with nocodazole are shown as a G2/M phase arrest control *p<0.05 by unpaired T-test. **e.** Cytogenetic analysis of the ALT+ *SMARCAL1* dependent osteosarcoma cell line, OS384 with adoxycycline inducible CRISPRi system against *SMARCAL1* vs. a gene desert control (Chr2.2). X axis shows the length of the doxycycline treatment for both the control and *SMARCAL1* KO cells; y axis represents the metaphase chromosome count. * p<0.05, ** p<0.005 **** p<0.0001 by unpaired T-test with Welch’s correction. **f.** Western immunoblotting confirming *SMARCAL1* repression upon doxycycline induction. GAPDH is used as a loading control.

Next, we asked whether *SMARCAL1*’s canonical role in replication fork stress responses explained its ALT-selective dependency. We perturbed *SMARCAL1* in ALT+ and non-ALT osteosarcoma models by CRISPR and measured markers of replication stress and checkpoint activation by immunoblotting (WB). After loss of *SMARCAL1*, we observed increased phosphorylation of RPA in both ALT+ and non-ALT models, consistent with SMARCAL1’s established role in replication stress in all cells.^53,54^ However, we observed a key divergence at the G2/M checkpoint. In ALT+, *SMARCAL1*-dependent cells, *SMARCAL1* loss increased CHK1 phosphorylation, whereas non-ALT model or *SMARCAL1* non-dependent models showed little or no change (**Fig. 6c**). Interestingly, this included the U2OS line, an ALT+ osteosarcoma cell line that does not depend on *SMARCAL1* (see **Discussion**). This pattern suggests that ALT+, *SMARCAL1* dependent cells engaged a stronger G2/M checkpoint response after *SMARCAL1* loss.

To quantify cell-cycle consequences without confounding toxicity from DNA double-strand breaks, we used an inducible CRISPRi system that silenced SMARCAL1 with catalytically-dead dCas9. We induced *SMARCAL1* loss in an ALT+ osteosarcoma model and performed flow cytometry after 14 days. We observed accumulation of cells in S phase and G2/M relative to controls (**Fig. 6d**). These data indicate that *SMARCAL1* suppression increased replication stress as expected and also caused a pronounced accumulation at the G2/M boundary in ALT+ dependent cells, consistent with defective progression through late cell cycle.

We next investigated the mechanism of cell death after SMARCAL1 depletion. By flow cytometry, we failed to visualize any significant annexin V-positive apoptosis signal using our CRISPRi system after SMARCAL1 depletion (**Extended Data Fig. 9a**). Furthermore, induction of DNA damage by γH_2_AX IF or WB was not markedly different between ALT+ and non-ALT models, similar to what has previously been shown by other groups (data not shown).^32,53,54^

These results suggested that checkpoint failure under replication stress might drive catastrophic outcomes. We therefore tested whether SMARCAL1 suppression promoted aneuploidy in ALT+ cells. We induced SMARCAL1 repression by CRISPRi in the ALT+ osteosarcoma line OS384 and performed spectral karyotyping at multiple time points up to 14 days. We observed significantly increased (*P*<0.05) chromosome counts after SMARCAL1 depletion relative to controls (**Fig. 6e**, **Extended Data Fig. 9b**) Take together, our results support a model in which *SMARCAL*1 loss destabilizes ALT telomere maintenance, increases C-circle production in *SMARCAL1*-dependent cells, and triggers a DNA damage checkpoint response that culminates in mitotic catastrophe and subsequent cell death.

## Discussion

ALT defines a clinically important subset of tumors that maintain telomeres without telomerase and are often associated with aggressive behavior.^8,9,12,69^ Despite long-standing interest in ALT as both a biomarker and a therapeutic opportunity, the field has lacked a practical way to call ALT at scale with high confidence. Experimental assays such as C-circles or telomere FISH provide strong evidence when applied carefully, but they require specialized expertise and do not fit easily into large discovery workflows. Computational approaches that quantify telomere content, variant repeats, fusions, or other WGS-derived features have helped, but most methods either rely on matched germline sequencing or struggle with batch effects, which limits their use in aggregated panels of cancer models.^15–17^ In this work, we addressed these constraints by developing ALTitude, which integrates telomere-relevant signals from WGS into a single model that does not require paired germline DNA and explicitly corrects technical variation. ALTitude matched orthogonal ALT assays with near-perfect accuracy in cell lines and enabled systematic annotation of ALT status across DepMap, a large public functional genomic and chemical screening database which includes genomic, epigenomic, and chemical response data in over 1,000 cancer cell lines.^19,20,70^ Thus, it will be available prospectively to support ongoing cancer discovery. To our knowledge, this is the first such tool available integrating ALT status with cancer genomic data in cancer cell lines.

The ability to annotate TMMs at scale means ALT can be used to stratify cell lines and link ALT to functional genomics, drug screens, and other omics data. Leveraging this new capability, we nominated *SMARCAL1* as a top ALT-associated dependency. Multiple studies have linked *SMARCAL1* to ALT biology in high-grade gliomas. In adult gliomas, investigators recently reported that *SMARCAL1* loss promoted ALT features.^45–47^ In contrast, in ALT+ gliomas that carry *ATRX* mutations, recent work has identified *SMARCAL1* as a selective dependency.^33^ These observations fit with the established role of SMARCAL1 in replication-stress responses, including at telomeres; SMARCAL1 functions as an annealing helicase at stalled replication forks, where it interacts with RPA-coated single-stranded DNA and promotes fork stabilization and restart.^53,54^ Though this function appears critical at telomeres, prior studies disagree on whether SMARCAL1 is exclusively important in ALT, or if SMARCAL1 is important more broadly at telomeres regardless of TMM.^32,71–74^ Consistent with its canonical role, *SMARCAL1* loss increased replication stress markers in both ALT+ and non-ALT cells. However, as shown in our RNA-seq and functional data, replication stress alone did not explain the selective dependency we observed. In ALT+, *SMARCAL1*-dependent models, *SMARCAL1* loss amplified ALT hallmarks, triggered stronger checkpoint signaling, and drove progressive genomic instability that culminated in aneuploidy and loss of viability over time. These findings support a model in which SMARCAL1 depletion triggers replication stress that ALT+ and non-ALT cells handle in fundamentally different ways. Non-ALT cells activate DNA damage response pathways, slow or arrest the cell cycle, and engage stress-response programs compatible with survival. ALT+ cells instead fail to mount these protective responses and continue proliferating under stress, leading to catastrophic genomic consequences and loss of viability. Our *in vivo* pooled screen in an ALT+ osteosarcoma cell line xenograft reinforced the relevance of this dependency in a disease context and prioritized *SMARCAL1* relative to other osteosarcoma-enriched liabilities that can be further explored in the future.

Though *ATRX* and *DAXX* have long been linked to ALT, the nature of *ATRX* “loss” varies widely across cancers and datasets.^48,75,76^ *ATRX* and *DAXX* are mutated at variable rates in different tumor types. *DAXX* alterations are most commonly associated with pancreatic neuroendocrine tumors, where they are highly concordant with ALT.^10^ Mutations or deletions in *ATRX* are found often in sarcomas and high-grade brain tumors, two tumor types with high rates of ALT.^10,40,48,65,77^ By re-calling and manually reviewing *ATRX* and *DAXX* events across DepMap, we found that ALT associated most strongly with complete loss of canonical ATRX function. Many *ATRX* alterations in non-ALT models appeared compatible with retention of a functional full-length allele, and those cases did not track with ALTitude-defined ALT status. This distinction matters for biomarker development: it suggests that simple presence or absence of any *ATRX* mutation will often misclassify ALT-relevant *ATRX* disruption, and it underscores the value of careful variant interpretation in large-scale dependency analyses.

A central result of this study is that the relationship between ALT and *SMARCAL1* dependency depends on the status of *ATRX*, particularly in certain tumor types like osteosarcoma. Our functional experiments support a compensatory relationship between ATRX and SMARCAL1 in a subset of ALT contexts. In osteosarcoma models, we demonstrate that altering the balance between these remodelers changed both fitness and ALT molecular readouts. These observations provide a mechanistic framework for recent work that reports that *SMARCAL1* loss can contribute to ALT phenotypes in some settings while acting as a selective dependency in others.^33,45–47^ One interpretation is that ALT+ cells tolerate loss of one of these activities, and in some genetic contexts, they even benefit from shifting the system toward a particular state, but they fail when they lose both. Additionally, our finding that *ATRX* itself represents a vulnerability in ALT when its function is preserved suggests an even more nuanced view of the roles of these homologous chromatin remodelers in creating and sustaining ALT. Our neuroblastoma data add further subtlety. IFF events in *ATRX* commonly remove the ADD domain, which binds heterochromatin marks and contributes to ATRX chromatin targeting.^43^ The inability of these truncations to phenocopy full-length ATRX in the context of *SMARCAL1* dependency raises the possibility that correct chromatin recruitment can also determine whether ATRX activity can substitute for SMARCAL1. This represents a testable hypothesis for future work, particularly to define the structural elements of ATRX that confer functional redundancy.

These findings also have therapeutic implications. Several features argue that *SMARCAL1* may offer a therapeutic window in *ATRX* loss, ALT+ cancers, in particular pediatric cancers like osteosarcoma. *SMARCAL1* encodes an enzyme with a defined catalytic core, and related SNF2 ATPases have increasingly become drug targets.^78^ In addition, *SMARCAL1* loss does not cause embryonic lethality in mouse models,^79^ which suggests tolerability of partial inhibition *in vivo*, although this observation does not strictly preclude on-target toxicities in proliferative tissues. The pediatric enrichment we observed is especially notable because high-risk pediatric sarcomas and *ATRX*-altered neuroblastoma often lack recurrent, actionable oncogenic drivers. In many such settings, a biomarker-linked synthetic lethal dependency may provide a more realistic route to targeted therapy than traditional driver inhibition.

Several limitations shape how broadly these results should be interpreted. First, ALTitude training relied on the subset of cell lines with orthogonally validated ALT status, and those models do not represent all tumor types evenly. Nonetheless, because ALTitude learned telomere sequence and structural features rather than lineage-specific expression programs, we expect it to generalize across diseases when high-quality WGS data are available. Cell lines remain imperfect models for tumors. Adaptation to long-term culture can alter telomere dynamics and stress responses, and some widely used ALT models behave atypically. For example, the osteosarcoma cell line U2OS,^80^ which has been cultured since the 1960s, expresses truncated ATRX, and is used in many fundamental mechanistic studies of SMARCAL1 and ALT+,^53,54^ does not show strong *SMARCAL1* dependency, highlighting that additional compensatory mechanisms can sustain ALT in certain backgrounds.

We report that SMARCAL1 rescues ATRX function in a subset of ALT. Though not tested here, this would presumably extend to other cancer lineages such as high-grade gliomas, for example, where *SMARCAL1* has been reported to be a critical survival gene in the *ATRX*-mutant ALT context. Based on our pan-cancer data, however, not all ALT+ cell lines with *SMARCAL1* dependency have apparent *ATRX* or *DAXX* genomic events. This is true, for example, in pediatric rhabdomyosarcoma, where only one out of four ALT+ lines dependent on *SMARCAL1* had loss of *ATRX*. Our data suggests the need for scrutiny of other roles for SMARCAL1 in ALT in these *ATRX* WT contexts. Finally, not all *SMARCAL1*-dependent lines are ALT+, indicating that *SMARCAL1* dependency can arise through alternative mechanisms in a minority of contexts, and it emphasizes the importance of using ALT status as a stratifier rather than assuming equivalence of *SMARCAL1* dependency to ALT. However, the role of *SMARCAL1* in replication stress response more generally, as well as recent literature demonstrating its regulation of innate immune recognition of non-ALT tumor tissues, suggests there is scope to target *SMARCAL1* more broadly beyond ALT in cancer.^64^

In summary, ALTitude provides a scalable and accurate framework for calling ALT from WGS without matched germline DNA and enables systematic integration of ALT status with functional genomics and pharmacology. Using this approach, we identified and validated *SMARCAL1* as a prominent ALT-associated dependency, enriched in pediatric ALT+ cancers and functionally linked in some contexts to *ATRX* status. These results define a tractable path for biomarker-guided target discovery in ALT and establish a foundation for developing therapeutic strategies tailored to ALT+ malignancies.

## Declaration of Interests

G.G. receives research funds from IBM, Pharmacyclics/Abbvie, Bayer, Genentech, Calico, Ultima Genomics, Inocras, Google, Kite, and Novartis and is also an inventor on patent applications filed by the Broad Institute including those related to MSMuTect, MSMutSig, POLYSOLVER, SignatureAnalyzer-GPU, MSEye, MinimuMM-seq, DLBclass, HapASeg, Tonly2, MSIDetect, TuFEst, and ABSOLUTE2. A full list of Dr. Getz’s publicly available patent applications can be requested from the Broad’s legal team at. He is a founder, consultant, and holds privately held equity in Scorpion Therapeutics; he is also a founder of and holds privately held equity in Predicta Biosciences; and holds privately held equity in Antares Therapeutics. K.S. previously received grant funding from Novartis and is on the SAB and has stock options with Auron Therapeutics. F.V. receives research support from the Dependency Map Consortium, Bristol Myers Squibb, Merck, Illumina, and AstraZeneca. F.V. has served as a consultant for GSK, and Riva Therapeutics. F.V. is a co-founder and equity holder in Jumble Therapeutics and an equity holder and consultant for Z Prime Therapeutics.

## Acknowledgements

We thank the following shared resource facilities at the St. Jude Comprehensive Cancer Center for their support of the studies in this manuscript: Cytogenetics, Flow Cytometry and Cell Sorting, and the Center for Advanced Genome Engineering. These resources and other work in this manuscript [in part] are supported by the American Lebanese-Syrian Associated Charities (ALSAC) and NCI P30-CA021765 (CWMR, Principal Investigator). We thank Drs. Jeffrey Klco and Ola Myklebost for contributions of cell line pellets for c-circle analysis. CWMR is supported by grants from the National Cancer Institute R01-CA-113794, R01-CA-273455, and R01-CA-172152. EB is supported by NIH R01-NS110837. KS received grant funding from the NCI R35-CA283977. PG is supported by an NIGMS R35-GM138293, an NCI R01-CA260060; NGRI K99/R00 R00-HG009679. This research was supported by the Office of the Assistant Secretary of Defense for Health Affairs, through the Peer Reviewed Cancer Research Program under Award No. HT9425-23-PRCRP-IA (LMG and EB). The content is solely the responsibility of the authors and does not necessarily represent the official views of the Department of Defense. We thank the Childhood Cancer Research Fund, the Rally Foundation for Childhood Cancer Research, the Sarcoma Alliance for Research through Collaboration, and the Damon Runyon Cancer Research Foundation for their support of these studies (LMG).

## ONLINE METHODS

### Ethics Statement

All research complied with ethical guidelines determined by St. Jude Children’s Research Hospital, Dana-Farber Cancer Institute, and the Broad Institute of Harvard and MIT. All animal studies were conducted at Dana-Farber Cancer Institute approved by the Institutional Animal Care and Use Committee under Animal Welfare Assurance number D16-00010 (A3023-01). No human studies were performed.

### Whole Genome Sequencing and Variant Analysis

Human genome assembly GRCh38 was used as a reference for mapping WGS and WES data using bwa (v0.7.12-r1039) with option “aln”. SNVs and indels were detected Bambino^81^ followed by a postprocess filtering^82,83^ including computational error suppression.^84,85^ Copy number variations (CNV) are detected by CONSERTING^86^ and structural variations (SV) detected by CREST. Candidate markers were evaluated for potential pathogenicity using PeCanPIE.^87^ All putative coding SNVs and indels and SVs were manually confirmed in Integrative Genomics Viewer (IGV).^88^ DNA-based mutations are further confirmed in RNAseq data whenever applicable to ensure the expression of the mutant allele. Biallelic *ATRX* loss was determined in samples with pathogenic ATRX variants that showed minimal wild-type full-length ATRX expression or evidence of biallelic inactivation of *ATRX* and no effect on fitness upon CRISPR-mediated *ATRX* knockout.

We used IGV to manually review the aligned sequence reads for every suspected variant allele identified for *ATRX*, *DAXX*, *SMARCAL1*, and *TERT* promoters across 936 cell lines total. To next resolve “complete loss of canonical ATRX function”, we additionally manually interrogated orthogonal data types, including RNA-seq and gene dependency scores for *ATRX* itself, allowing us to distinguish true ATRX-loss-of-function lines from cell lines with residual ATRX function. Because *ATRX* is on the X chromosome, we also considered donor sex and the possibility of functional duplication of the mutant or wild-type X.

### ALTitude method for predicting ALT in cancer cell lines

#### Telomere Feature Extraction

To characterize telomere biology, we applied a standardized computational pipeline to GRCh38-aligned WGS data. Average telomere length was estimated using *Telseq*,^15^ which quantifies telomeric reads containing a defined number of TTAGGG hexameric repeats in 150 bp reads and normalizes counts by the number of reads with a GC content between 48–52%. Telomeric fusion events were detected using *TelFusDetector*,^16^ and telomeric variant repeats (TVRs) were identified using *TelomereHunter*.^17^ Neo-telomere structures were inferred from whole genome sequencing data using *Telfuse*.^18^

#### Batch Effect Correction

To mitigate technical confounding due to sequencing batch effects, telomere features were normalized using a principal component regression approach. Specifically, Singular Value Decomposition (SVD) was applied to the centered mosdepth sequencing coverage matrix (genomic regions × samples) to derive principal components capturing technical variation. Each telomere feature was then log-transformed (with feature-specific offsets to handle zero values) and regressed against the first 10 depth-derived principal components using ordinary least squares regression. The residuals from these regressions, representing telomere signal independent of sequencing depth variation, were retained as the batch-corrected normalized features, previously described by Taub et al.^89^ Sequencing depth in 1kb bins was calculated using mosdepth.^90^

#### Machine learning Model

To develop the machine learning predictive model, we compiled a training dataset of 134 pediatric and adult cancer cell lines with ALT informative assays available such as C-circle assay, APB FISH, and TRAP assay. After quality filtering, 82 cell lines with complete data were retained for model development. ALT status labels were assigned using unsupervised hierarchical clustering based on the batch-corrected telomere features described above: telomere content, telomere fusion rate, neo-telomere frequency, TERT expression, and telomeric variant repeat (TVR) content (represented as 49 individual hexamer variant frequencies, yielding 53 total features).

We evaluated multiple machine learning approaches including a stacked ensemble combining gradient boosting and tree-based classifiers with a logistic regression meta-learner. The base models included Logistic Regression, Random Forest, XGBoost, CatBoost, LightGBM, ExtraTrees, Histogram-based Gradient Boosting, AdaBoost, and TabPFN2, all using default hyperparameters. Missing values were handled natively by gradient boosting models (XGBoost, CatBoost, LightGBM, HistGB) or imputed using mean imputation for other classifiers. Individual base models were trained using leave-one-out cross-validation on the training set (N=57, 70% of data), with a held-out test set (N=25, 30%) for final performance evaluation. The logistic regression meta-learner was trained on out-of-fold predictions from base models and validated using k-fold cross-validation, achieving an AUROC of 1.0. TabPFN2, a neural network-based tabular classifier, achieved equivalent performance (AUROC = 1.0) to the stacked ensemble while providing well-calibrated probability estimates. We therefore selected TabPFN2 as the final classifier and applied a probability threshold of >0.6 to classify cell lines as ALT+.

#### Feature ablation analysis

To assess model robustness and identify minimal predictive feature sets, we performed a comprehensive ablation study across 20 feature configurations and 9 model architectures (180 total experiments). Feature sets ranged from single features (e.g., neo-telomere count alone) to the full 53-feature model, including intermediate combinations (e.g., neo-telomere + fusion rate, core 4 features). For each configuration, we evaluated both cross-validated AUROC and Jaccard similarity of ALT+ predictions compared to the full ensemble model, ensuring that high discriminative performance translated to consistent sample-level classifications.

### Comparison to RNA-based classifier

To benchmark ALTitude against existing methods, we compared performance to a recently published RNA-based ALT classifier⁵ on 82 overlapping samples with matched WGS and RNA-seq data. We reimplemented the published approach using 5-fold cross-validation with feature selection performed within each training fold to prevent data leakage, and evaluated both methods on the same 82 samples under identical cross-validation splits. For deployment, TabPFN2 was retrained on 100% of the labeled data (N=82), and ALT status was predicted for 825 additional DepMap cell lines (906 total with WGS-derived features).

Feature stability was assessed through an ablation analysis across 20 feature configurations. All analyses used a fixed random seed for reproducibility. The complete pipeline was containerized using Singularity and implemented in Workflow Description Language (WDL) for reproducible execution across computational environments. ALT calls will be made available from the PedDep portal. All code is available on GitHub/Zenodo. Differential dependencies were identified using linear modeling with empirical Bayes moderation (limma). Statistical models and visualizations on CRISPR-KO data were generated with R (v4.1.2).

### Structural alignment

The ATP-binding subdomains of ATRX (residues 1,581-1,768) and SMARCAL1 (residues 445-600) were extracted and aligned using ClustalW via the msa package (v1.28.0) in R (v4.3.0). Sequence conservation was calculated at each aligned position: identical residues were scored as conserved, chemically similar substitutions as partially conserved, and gaps or dissimilar residues as non-conserved. Percent sequence identity was calculated as the number of identical positions divided by the total number of aligned positions excluding gaps.

AlphaFold-predicted structures for ATRX (AF-P46100-F1) and SMARCAL1 (AF-Q9NZC9-F1) were obtained from the AlphaFold Protein Structure Database. The ATP-binding subdomains were extracted and structurally aligned using PyMOL (v2.5). Structural superposition was performed using the align command with 5 refinement cycles. Root-mean-square deviation (RMSD) was calculated over Cα atoms of aligned residues. The structural superposition image was rendered in PyMOL with ray tracing at 300 DPI. Conservation tracks display aligned residue positions colored by conservation status: identical (red), similar (orange), or gap/dissimilar (grey).

### Sterile tissue culture

All cell lines were cultured in humidified incubators at 37°C with 5% CO_2_. The neuroblastoma cell lines CHLA90, SKNMM, SKNFI and the rhabdomyosarcoma cell lines RH4, RH28, RH30, RH41, RHJT were provided by Dr. Adam Durbin (St. Jude Children’s Research Hospital). The neuroblastoma lines COGN512 and CHLA119 were provided by Dr. C. Patrick Reynolds (Children’s Oncology Group cell line repository at Texas Tech University Health Sciences Center). The osteosarcoma cell lines, G292, MG63, CAL72 and U2OS were gifts from Dr Kimberly Stegmaier (Dana-Farber Cancer Institute). The PDX-derived OS525 and OS384 cell lines were a gift from Dr Alejandro Sweet-Cordero (UCSF Benioff Children’s Hospital). The NY cell line was purchased from XenoTech,LLC (Kansas City, KS). Cells were maintained in growth media listed below supplemented with 1% penicillin-streptomycin (Thermo Fisher, Waltham, MA).

| Cell line | Growth medium |
| --- | --- |
| CHLA90 | DMEM, 20% FBS |
| SKNMM | RPMI1640, 10% FBS |
| SKNFI | DMEM, 10% FBS |
| COGN512 | IMDM, 20% FBS, 1% ITS |
| CHLA119 | IMDM, 20% FBS, 1% ITS |
| OS525 | DMEM, 10% BGS |
| OS384 | DMEM, 10% BGS |
| NY | EMEM, 10% FBS |
| G292 | McCoy's 5A, 10% FBS |
| MG63 | EMEM, 10% FBS |
| U2OS | McCoy's 5A, 10% FBS |
| CAL72 | DMEM, 10% FBS, 1% ITS |
| SJSA1 | RPMI1640, 10% FBS |
| RH4 | DMEM, 10% FBS |
| RH28 | DMEM, 10% FBS |
| RH30 | DMEM, 10% FBS |
| RH41 | DMEM, 10% FBS |
| RHJT | DMEM, 10% FBS |

### Lentivirus production and CRISPR knockout

For lentivirus generation HEK 293T cells (ATCC, Manasass, VA #CRL-3216) were plated in 10-cm tissue culture treated dishes (Corning, Glendale, AZ #430293) at 3.5X10⁶ cells/plate density. 24 hours later 3μg of the lentiviral construct and 3μg of psPAX2 and pMD2.G (Addgene, Watertown MA#12260 and #12259) packaging plasmids were added with 18 μL XtremeGene360 transfection reagent (EMDMillipore, Burlington, MA #XTG360-RO). After 48 hours, viral particles were collected and filtered through a 0.45 µM membrane (EDM Millipore, Burlington, MA #SEIM003M00).

Lentiviral infections were carried out in 10-cm dishes (Corning, Glendale, AZ #430293) using 1X10⁶ cells/plate. After removal of the normal growth medium, cells were covered with 2 mL of virus, and 1.6µL of 10mg/mL polybrene transfection reagent (EMD Millipore, Burlington, MA #TR-1003-6). After 4 hours growth media was added without virus removal. Selection started 48 hours post infection and cells were harvested for further experiments 2-6 days post-selection.

### CRISPR cloning

Gene-specific guide RNA oligonucleotides were purchased from Integrated DNA Technologies (Coralville, Iowa) using the Avana library sequences (available at www.depmap.org).^91^ The small guide RNAs (sgRNAs) were cloned into the following Lentiviral vectors by previously described methods:^92,93^ pLCKO2 #1255180, LentiCRISPRv2puro #98290 and lentiCRISPRv2blast #98293 (Addgene, Watertown,MA). All sequences were confirmed by next generation sequencing (NGS) (Plasmidsaurus Inc. Eugene, OR or Genewiz Azenta Life Sciences, South Plainfield, NJ). Endofree DNA was prepared using the ZymoPURE II maxiprep kit (Zymo Research, Irvine, CA #4202).

For inducible CRISPR/Cas9 experiments sgRNAs were cloned into the FgH1tUTG Lentiviral vector (Addgene, Watertown MA #70183). These constructs were transduced into mammalian cell lines that constitutively express the Cas9 endonuclease from the pLX311-Cas9 plasmid (Addgene, Watertown MA #118018).

CRISPR guide sequences:

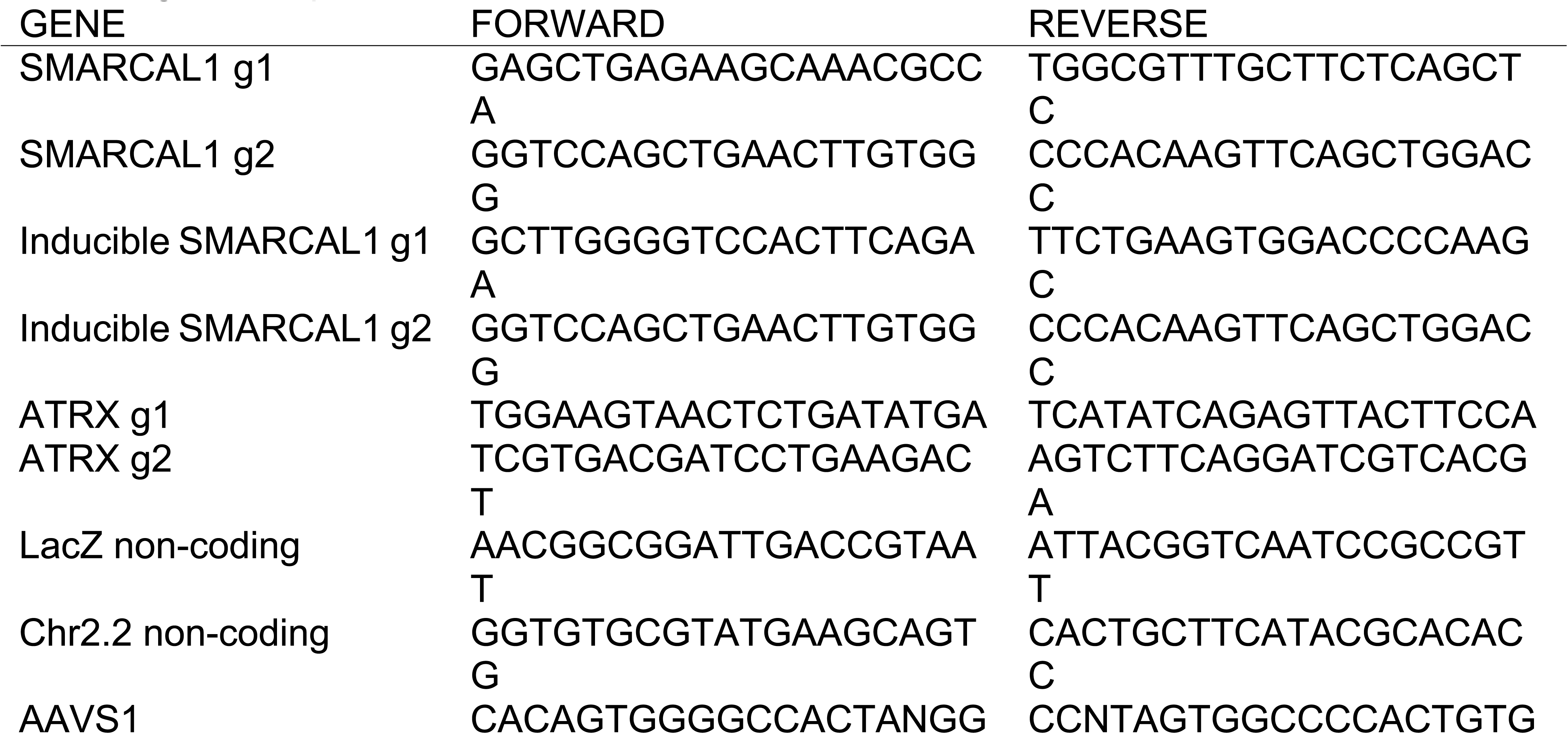

### dTAG construct cloning and validation

For rapid target protein degradation, the dTAG system was used as previously described.^38^ Briefly, exogenous expression of a target protein fused at the carboxy-terminal, to a 2xHA tag as well as the FKBP12^F36V^ protein fragment, targets the protein for degradation in the proteosome upon treatment with an E3 ubiquitin ligase-recruiting small molecule. The target gene *SMARCAL1* cDNA was developed as a gBlock (IDT, Coralville, IA) with silent mutations introduced into all PAM sites, rendering it resistant to CRISPR KO. This was then cloned into the Lentiviral vector pLEX_305-C-dTAG (Addgene, Watertown, MA #91798) via Gateway Cloning (Thermo Fisher, Waltham, MA) according to the manufacturer instructions. Mammalian cell lines then were infected by the lentiviral infection method described above. Simultaneously, KO of endogenous *SMARCAL1* via CRISPR was performed (see above) such that the exogenous, tagged *SMARCAL1* was the predominant SMARCAL1 protein expressed. This was confirmed by Western immunoblotting. Cells expressing the exogenous, C-terminally tagged proteins, were treated with various concentrations of dTAG^V^-1 (Tocris, Bristol, UK #6914). Degradation was monitored at various time points by western immunoblotting.

#### Overexpression plasmid constructs

The SMARCAL1 cDNA sequence was cloned into the mammalian lentiviral expression vector pLEX_307 (Addgene, Watertown, MA #41392) by Gateway technology. As a control, pLEX_307 expressing the wt GFP ORF was constructed. The ATRX and EGFP overexpression plasmids were generated by VectorBuilder (https://www.vectorbuilder.com). The VB240919-1575xjq plasmid expresses the full length ATRX ORF and the VB240919-1581jqn vector expresses the EGFP ORF both modified with an HA tag and driven by a CMV promoter. The plasmid sequences are available upon request. The overexpression plasmids were used for Lentivirus production as described above.

### Western immunoblotting

Cell extracts were prepared in 1X Lysis buffer (Cell Signaling Technologies, Danvers, MA #9803) supplemented with PhosSTOP (#04906837001) phosphatase and cOmplete Mini (#11836153001) protease inhibitor tablets (Roche-Sigma-Aldrich, St. Louis, MO). Protein concentrations were determined using the Bradford reagent (BioRad, Hercules, CA #5000205) according to the manufacturer’s protocol. Protein samples were run on precast gels (Invitrogen-Thermo Fisher, Waltham, MA), transferred to Immobilon FL membranes (EDM Millipore, Burlington, MA #IPFL 00010) and detected by chemiluminescence using the Super-Signal West Dura substrate (Thermo Fisher, Waltham, MA #34076) by the manufacturer recommendation. Imaging was done using the Li-COR Odyssey Fc (LI-COR, Lincoln, NE) or on the BioRad ChemiDoc Touch imaging systems (BioRad, Hercules, CA #12003154). Antibodies used are listed below.

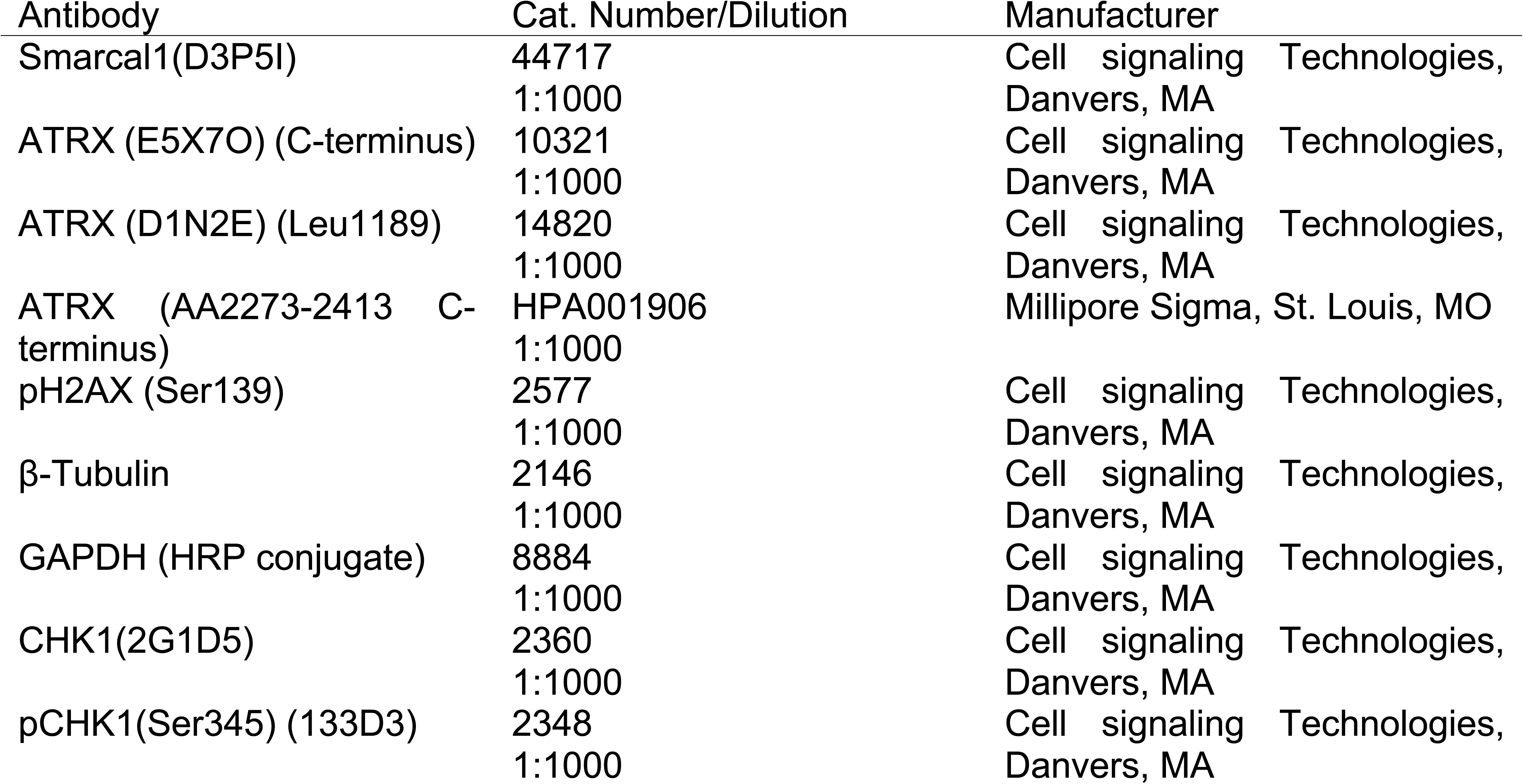
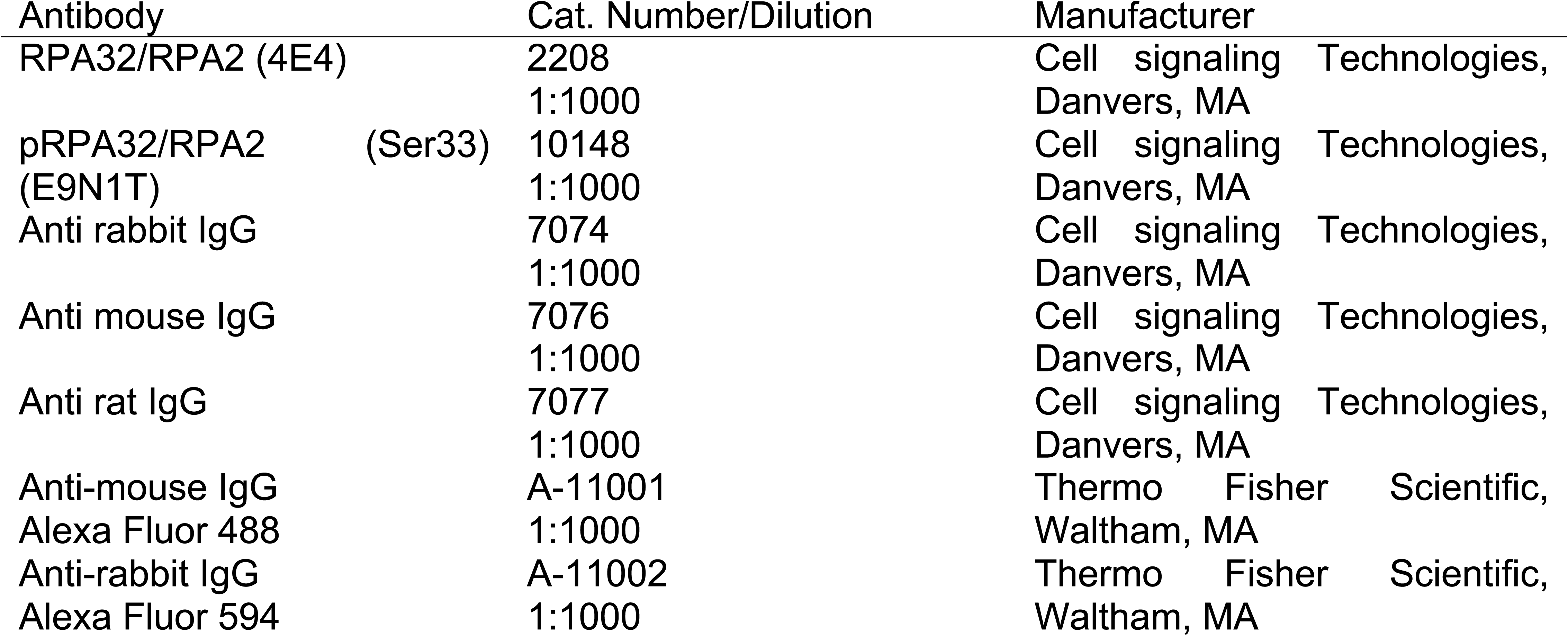

### Animal procurement

Animal care and use was conducted in accordance with Dana-Farber Cancer Institute institutional guidelines and relevant regulatory standards. All studies were approved by the Institutional Animal Care and Use Committee under Animal Welfare Assurance number D16-00010 (A3023-01).

### *In vivo* pooled screen and parallel *in vitro* screen methods

Using DepMap Release Version 19Q1 (www.depmap.org), 108 genes of interest were chosen enriched for dependency in osteosarcoma. The genes chosen were enriched for dependency in osteosarcoma vs. all other cell lines with effect size < -0.1, p value < 0.05, with dependency (Ceres score <-0.5) in at least 2 out of 6 osteosarcoma cell lines. Genes that were targets in OS but considered pan-essential by CRISPR screening were included (13). sgRNAs were chosen from the Broad Institute Avana and Brunello libraries, including the top two performing Avana and top two Brunello guides (or tiered Brunello guides if top 1-2 Brunello guides were identical to Avana). In addition, 354 intronic controls for copy number/cutting toxicity were chosen using the GRCh38 reference genome. 600 cutting/intergenic controls were designed to target “deserts of function” in the genome. Guides were included that were not near an annotated coding sequence and then selected to the top sgRNAs in terms of specificity (i.e., only had a single hit in a “desert of function” plus have minimal predicted off-target effects when looking at 1-,2-, and 3-base mismatches), then gave preference to those with high predicted efficacy. Additionally, 200 non-targeting controls were chosen, as well as 10 additional positive controls. After synthesis of the barcoded library and QC, high-titer lentivirus was generated and infected into Cal72 cells with constitutive Cas9 expression using spin infection. After one day of infection and three days of selection in puromycin, cells were counted and injected subcutaneously into the flanks of 8-week-old NOD.Cg-Prkdcscid Il2rgtm1Wjl/SzJ (NSG) mice (Jackson Laboratory, Bar Harbor, ME). 3×10^6^ cells were injected per mouse, and cells were diluted in PBS. 15 mice were injected in total. In parallel, an *in vitro* screen was completed over 25 days (3 replicates) for parallel confirmation of sequencing barcodes and hits. Mice were taken down at 11 days (4 mice) and 25 days (4 mice), with cells collected for sequencing also from the parallel *in vitro* screen replicates. DNA was extracted and sequenced using Illumina NGS technology.

The gene-level dependencies from pooled CRISPR screens were quantified based on STARS v1.3, a computational tool developed by the Broad Institute’s Genetic Perturbation Platform (https://portals.broadinstitute.org/gpp/public/). For the pooled screen, STARS was applied with the user-defined threshold 30%. Significant gene dependency hits were identified based on the cut-offs STARS dependency score ≤ -1.5 and p-value ≤ 0.10.

### Cell viability assays

Cell viability was determined by the CellTiter-Glo® (CTG) Luminescent Assay (Promega, Medison, WI #G7573.) Generally, cells were seeded in growth medium to 384 well plates (Corrning, Glendale, AZ #3570) at a concentration of 2.5X10^3^ cells/mL, in a volume of 50 μL/well, in 4-8 replicates/condition. Viability was monitored up to 12 days. Cells were incubated 15 min at RT with 10 μL/well CTG reagent and luminescence was measured by a VANTAstar microplate reader (BMG Labtech Ortenberg, Germany). Growth curves were generated using the GraphPad Prism 9 software.

Cell viability was also assessed using the methylcellulose soft agar colony forming assay (StemCell Technologies, Cambridge, MA #03814). Generally, 2.25 mL methylcellulose was aliquoted into 14mL round bottom tubes and mixed with 0.25mL of 1X, 1.5X or 2X10^5^/ml cells. 0.5mL of cell suspension was aliquoted in triplicates to 24 well plates and incubated at 37°C for 9-11 days. Colonies were visualized after 4 hours of incubation with 100μL 2.5 mg/mL MTT per well (MilliporeSigma, Burlington, MA #475989). High resolution pictures were taken with the BioRad Chemidoc MP imaging system (BioRad, Hercules, CA #12003154).

### IncuCyte® Live-cell imaging

Proliferation assays were run and analyzed by an IncuCyte® S3 instrument (Sartorius, Gottingen, Germany.) Experiments were set up in 96 well plates (Corrning, Glendale, AZ #3904) with cell densities of 5-20X10³ cells /well in 4 replicates/condition. High resolution bright field images were taken at 4 different areas of each well every 4 hours for 5-7 days. Proliferation metrics were determined and analyzed by the IncuCyte® classic confluence analysis software.

### Cell cycle analysis

For cell cycle analysis, 5X10⁵ cells (floating and adherent) were used. After a DPBS (Corning,Glendale AZ #21-031-CM) wash, cells were incubated for 15 minutes at RT with a fixable viability dye, Ghost Dye Red 780 (Cytek Bioscienses Bethesda, MD #13-0865-100), followed by a wash in staining media (DPBS+1% FBS). Cells were fixed (100μL per sample, 15 minutes at RT) and permeabilized (100μl per sample,15 minutes at RT) using the Click-iT Plus EdU Alexa Fluor 488 Flow cytometry assay Kit’s (Invitrogen-Thermo Fisher, Waltham, MA #C10632) fixative and permeabilization buffer. Finally,cells were resuspended in 0.5mL permeabilization buffer and 1 drop of FxCycle Violet Ready Flow Reagent (Thermo Fisher, Waltham, MA # R37166), incubated 30 minutes at RT away from light before Flow Cytometry using the LSRFortessa cell analyzer (BD Biosciences, Franklin Lakes, NJ) and the FlowJo software.

### Apoptosis flow cytometry

2.5X10⁵ cells were washed in DPBS (Corning, Glendale AZ #21-031-CM) and resuspended in 100μL Annexin-V mix, containing 85μL 1X Annexin V Binding Buffer (BD Biosciences, Franklin Lakes, NJ #556454), 5μL Annexin V-APC (Cytek Biosciences, Bethesda, MD #20-6409-T025) and 10μL 1μg/mL Dapi stain (Thermo Fisher, Waltham, MA # P36931) After 15 minutes incubation at RT in the dark an additional 150μL cold Annexin V Binding Buffer was added to each sample. Flow Cytometry was performed on the LSRFortessa cell analyzer (BD biosciences, Franklin Lakes, NJ) and analyzed by the FlowJo software.

### Spectral Karyotype Analysis (SKY)

Actively dividing cells were treated with colcemid (Thermo Fisher, Waltham, MA. #15212-012) for 4hours. Spectral karyotyping was performed as previously described^94^ using a human SkyPaint DNA kit and the Concentrated Antibody Detection kit (Applied Spectral Imaging, Carlsbad, CA #FPRPR0028 and #FPRPR0033) according to the manufacturer protocol.

### C-Circle Assays

The C-circle assay was performed as previously described, with isothermal rolling-circle amplification carried out on an Applied Biosystems MiniAmp Thermal Cycler at 30 °C for 8 h, 65 °C for 20 min, and held at 4 °C for up to 24 h, using 32 ng template DNA, 2 µL BSA (2 µg/µL), 2 µL 1% Tween-20, 0.8 µL 100 µM DTT, 2 µL 10 mM dNTPs (NEB, Ipswitch, MA N0447L), 2 µL Φ29 buffer, 0.8 µL Φ29 DNA polymerase (NEB, M0269L), and nuclease-free water to 20 µL total volume, with corresponding no-Φ29 controls prepared identically except substituting nuclease-free water for Φ29 polymerase; all reactions were subsequently diluted to a final volume of 40 µL with nuclease-free water.

To detect amplification of telomeric repeats from C-circles, real-time PCR was performed on an Applied Biosystems QuantStudio 7 system using a MicroAmp Optical 96-well plate, with amplification of telomeric DNA (Forward primer: 5′-CGGTTTGTTTGGGTTTGGGTTTGGGTTTGGGTTTGGGTT-3′; Reverse primer: 5′-GGCTTGCCTTACCCTTACCCTTACCCTTACCCTTACCCT-3′) and VAV2 DNA (Forward primer: 5′-TGGGCATGACTGAAGATGAC-3′; Reverse primer: 5′-ATCTGCCCTCACCTTCTCAA-3′) (IDT, Coralville, IA) carried out in 25 µL reactions containing 5 µL of diluted isothermal product, 12.5 µL QuantiTect SYBR Green PCR Master Mix (Qiagen, Hilden, Germany 204445), 1 µL 100 µM DTT, 0.5 µL DMSO, 1 µL nuclease-free water, and 2.5 µL of each primer (1 µM F and 9 µM R for telomeres, or 4 µM F and R for VAV2), under cycling conditions of 95 °C for 15 min followed by 33 cycles of 95 °C for 15 s and 56 °C for 2 min for telomeres, or 40 cycles of 95 °C for 15 s, 57 °C for 30 s, and 72 °C for 1 min for VAV2, with CHLA90 and CHLA20 serving as positive and negative controls, respectively; all Φ29 and no-Φ29 reactions for telomeres and VAV2 were run in triplicate on the same plate to minimize inter-run variability, and samples were deemed CC⁺ if normalized values were ≥5 arbitrary units (AU) relative to CHLA90.

### RNA-seq

Total RNA was isolated from triplicate samples using the RNeasy Mini kit (Qiagen, Germantown, MD #74104). RNA was quantified using the Quant-iT RiboGreen RNA assay (ThermoFisher Waltham, MA # R11491) and quality checked by the 2100 Bioanalyzer RNA 6000 Nano assay (Agilent Santa Clara, CA #5067-1511) or 4200 TapeStation High Sensitivity RNA ScreenTape assay (Agilent Santa Clara, CA #G2991BA, #5067-5579) prior to library generation. Libraries were prepared from total RNA with the Illumina Stranded Total RNA Library Prep Kit according to the manufacturer’s instructions (Illumina,San Diego, CA # 20040529). Libraries were analyzed for insert size distribution using the 2100 BioAnalyzer High Sensitivity kit (Agilent Santa Clara, CA #5067-4626), 4200 TapeStation D1000 ScreenTape assay (Agilent #5067-5582), or 5300 Fragment Analyzer NGS fragment kit (Agilent #M5311AA). Libraries were quantified using the Quant-iT PicoGreen ds DNA assay (ThermoFisher Waltham, MA #P11496) or by low pass sequencing with a MiSeq nano kit (Illumina San Diego, CA # MS-103-1003). Paired end 100 cycle sequencing was performed on a NovaSeq X Plus (Illumina San Diego, CA # 20084804).

RNA-seq data from *SMARCAL1* CRISPR KO experiments were analyzed using a consistent methodology for both cell lines. For G292 (4 KO vs. 3 control, 72-hour timepoint), FPKM values were obtained from a combined RSEM output table. For Huh7 (6 KO vs. 5 control), FPKM values were obtained from GSE221372 (note: the Control_0 sample was excluded due to file formatting issues resulting in missing data). Differential expression was performed using limma on log2(FPKM+1) transformed values. Gene set enrichment analysis was performed using the fgsea R package with 10,000 permutations, ranking genes by limma moderated t-statistic. ALT gene signatures were derived from DepMap differential expression analysis comparing ALT+ (n = 39) and non-ALT (n = 362) cell lines. To reduce confounding by tissue-of-origin, the analysis was restricted to lineages containing at least one ALT+ model (398 cell lines total), and tissue substructure was accounted for by including principal components derived from the expression matrix as covariates. Differentially expressed genes were identified using relaxed thresholds (|log2FC| > 0.25, FDR < 0.2) and intersected with curated telomere gene sets, yielding 84 ALT-upregulated and 30 ALT-downregulated genes.

### Data Availability

Whole-genome sequencing data are available through dbGaP (accession: phs003444.v3.p1). CRISPR dependency and gene expression data were obtained from the Depmap Portal (PedDep/DepMap).

RNA sequencing data for Huh7 *SMARCAL1* wildtype and knock out cells are available from the Gene Expression Omnibus (GEO) under accession GSE221372. RNA sequencing data for G292 cells were generated in-house and are available upon request to the corresponding author.

### Code Availability

Custom code for telomere feature extraction (TabALT) is available at https://github.com/declan93/TabALT. The ALTitude machine learning pipeline for predicting ALT status from whole-genome sequencing data is available at https://github.com/declan93/ALTitude.

### Statistics and reproducibility

All statistics are as specified in the text, figures, and/or figure legends. Statistical analyses were performed using R (v4.3.3), Microsoft Excel, and GraphPad Prism. Figure panels displaying data from experiments with n=1 include Figs. 2, 5, and 6. All other figure panels display data from experiments with at least n = 2.

### Declaration of generative AI and AI-assisted technologies in the writing process

During the preparation of this work the author(s) used Claude code in order to refactor and condense code and to edit certain sections of the manuscript writing for clarity. After using this tool/service, the author(s) reviewed and edited the content as needed and take(s) full responsibility for the content of the published article.

## Extended Data Figures

**Extended Data Fig. 1.**
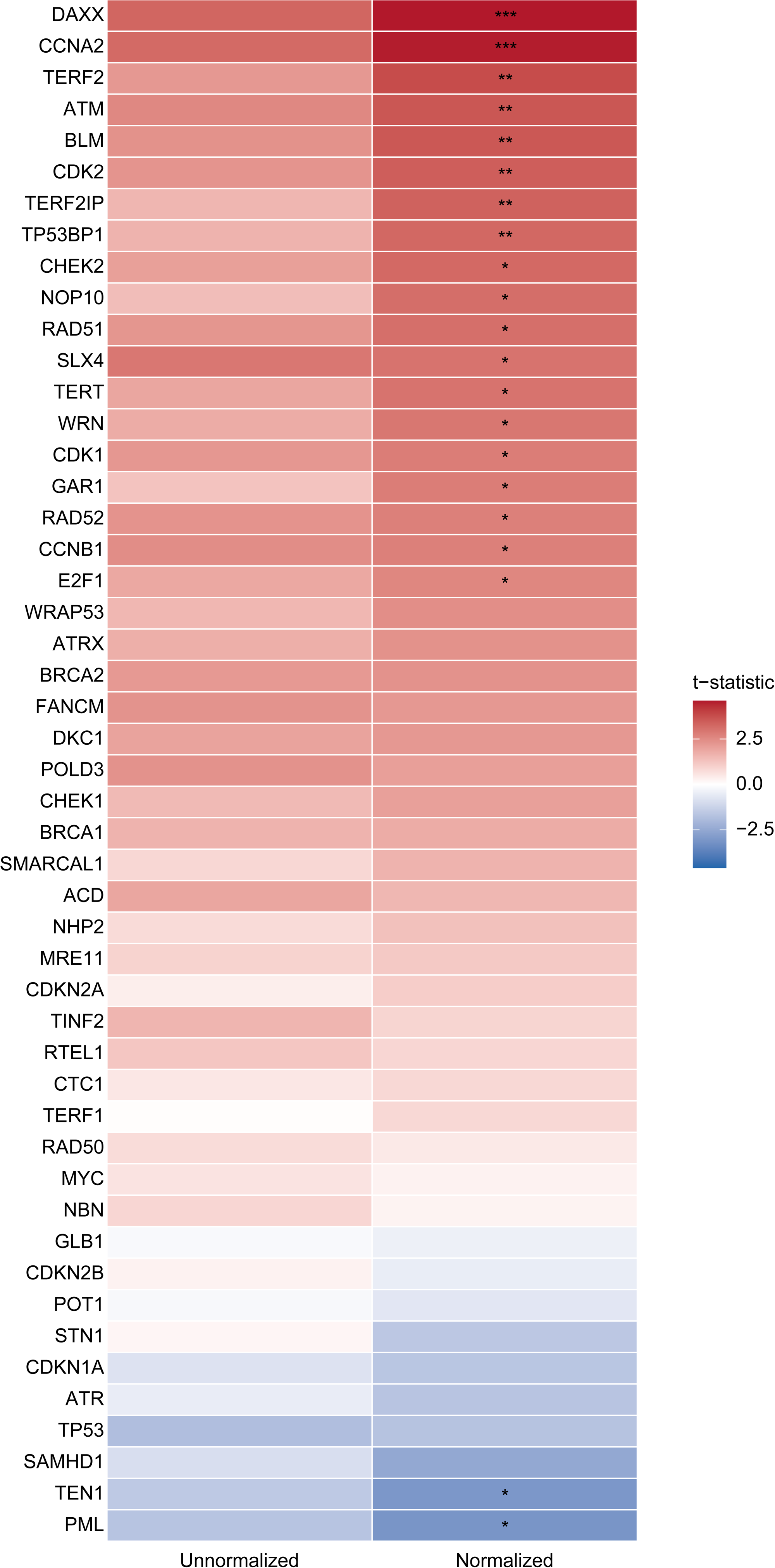
Normalized vs un-normalized telomere gene dependency enrichment. Gene expression in telomere-relevant genes as a function of telomere length. Heatmap displaying t-statistics from limma linear models testing the association between gene expression (TPM) and telomere content (TelSeq) for 49 candidate telomere-relevant genes. Left column shows results from unnormalized TelSeq values; right column shows results after TelSeq normalization (see Methods). Genes are ordered by normalized t-statistic. Positive values (red) indicate genes with increased expression in cell lines with longer telomeres; negative values (blue) indicate genes with decreased expressionin cell lines with longer telomeres. *p_adj < 0.05, **p_adj < 0.01, ***p_adj < 0.001 by limma moderated t-test with Benjamini-Hochberg correction.

**Extended Data Fig. 2.**
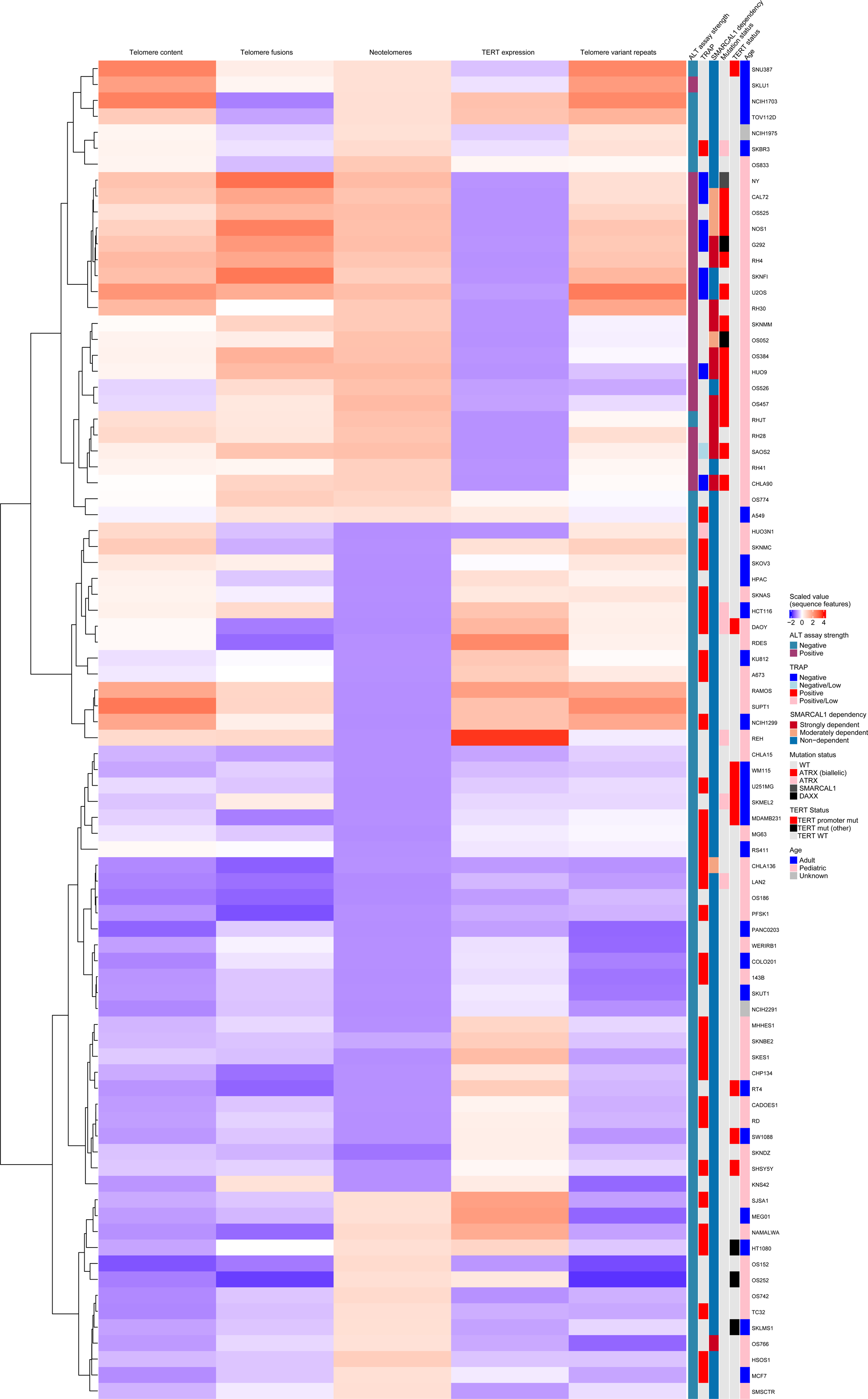
Training dataset heatmap. Telomere maintenance features across assay-confirmed cell lines. Heatmap of z-scaled telomere sequence features (telomere content, telomere fusions, neotelomeres, *TERT* expression, and telomere variant repeats) derived from whole-genome sequencing data across cell lines with experimentally confirmed ALT status. Rows (cell lines) were hierarchically clustered using Canberra distance with Ward’s D2 linkage. Columns represent individual telomere features. Row annotations indicate ALT assay strength (C-circle/APB status), TRAP assay results, *SMARCAL1* dependency classification (strongly dependent: Chronos score ≤ −0.5, moderately dependent: Chronos score ≤ −0.35, non-dependent: Chronos score > −0.35), mutation status (*ATRX* biallelic loss, *ATRX* other, *SMARCAL1*, *DAXX*, or wild-type), *TERT* status (promoter mutation, other *TERT* mutation, or wild-type), and age category (adult, pediatric, or unknown).

**Extended Data Fig 3.**
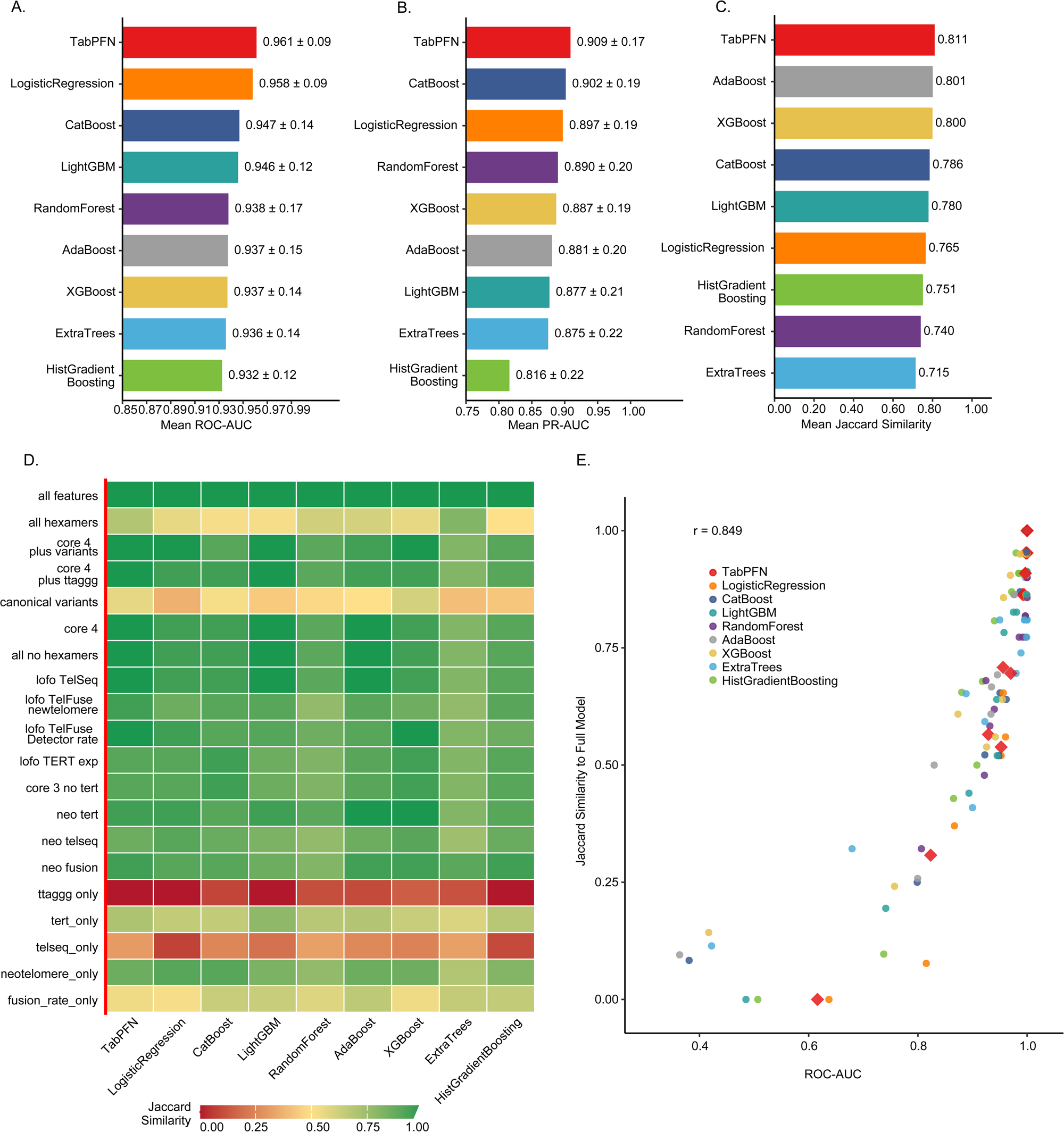
Ablation data. **a.** Mean ROC-AUC performance (± s.d.) for each machine learning model across 20 feature configurations. TabPFN2 achieved the highest mean ROC-AUC (0.9615), indicating superior stability across feature subsets. b. Mean PR-AUC performance (± s.d.) for each machine learning model across 20 feature configurations. c Mean Jaccard similarity to the reference full model (TabPFN with all 53 features) for each model type, averaged across all feature configurations. Higher values indicate more consistent predictions with the full model regardless of feature subset used. **d.** Heatmap of Jaccard similarity between each model–feature set combination and the reference full model predictions. Rows representfeature configurations ordered by complexity: hexamer-only features (top), single telomere features, minimal combinations (2–3 features), core features, combined sets, and leave-one-feature-out (LOFO) configurations (bottom). Columns represent models. Green indicates high agreement (Jaccard = 1.0); red indicates low agreement. TTAGGG hexamer alone shows universally poor agreement across all models, while core feature sets (Core 4, Core 4 + TTAGGG, all features) achieve near-perfect agreement for most models. **d.** Relationship between cross-validated ROC-AUC and Jaccard similarity to the full model across all 180 experiments. Each point represents one model trained on one feature configuration. TabPFN2 (red diamonds) clusters in the upper right, demonstrating both high performance and high prediction consistency. Pearson correlation r = 0.858.

**Extended Data Fig 4.**
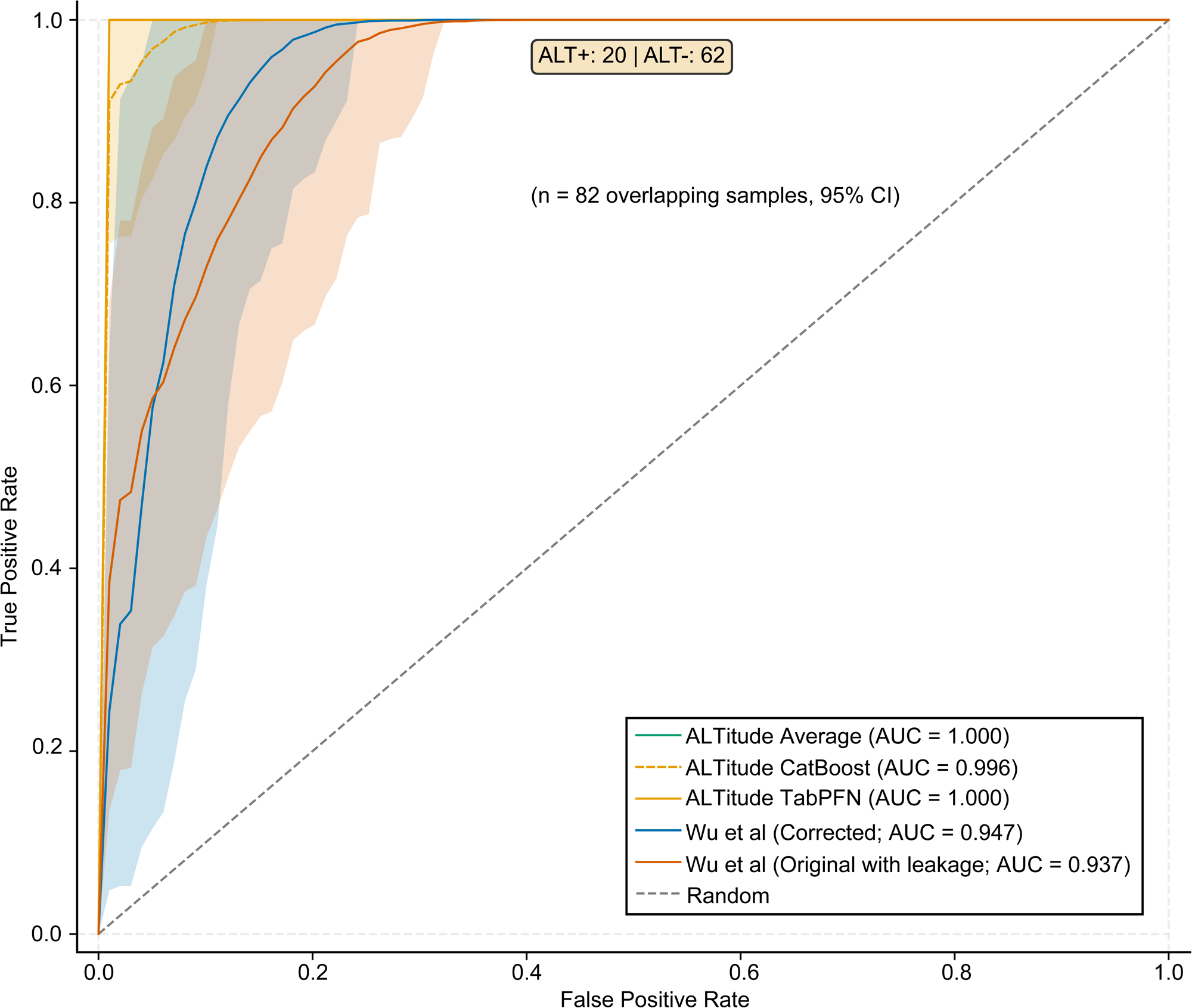
Comparative ROC curves. Receiver operating characteristic (ROC) curves comparing ALTitude models with the *Wu et al.* method on 82 overlapping cell lines (20 ALT+, 62 ALT−). Solid lines show ALTitude models: Ensemble (green), CatBoost (yellow dashed), and TabPFN2 (yellow), all achieving AUCs of 1, 0.996 & 1 respectively. Methods developed by *Wu et al.* are evaluated under two conditions: the original implementation containing original data leakage (orange dashed, AUC = 0.93758) and after correction for leakage (dark blue, AUC = 0.94733). Shaded regions indicate 95% confidence intervals from bootstrapping. The diagonal grey dashed line represents random classifier performance (AUC = 0.5). ALTitude models achieve perfect discrimination on this subset, outperforming the corrected *Wu et al.* method.

**Extended Data Fig. 5:**
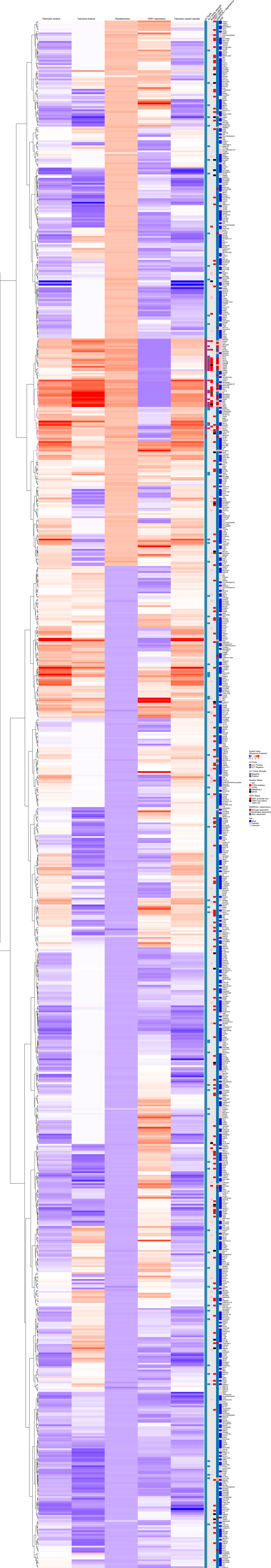
Full heatmap. Heatmap of z-scaled telomere sequence features (telomere content, telomere fusions, neotelomeres, TERT expression, and telomere variant repeats) derived from whole-genome sequencing data across all cell lines in the dataset. Rows (cell lines) were hierarchically clustered using Canberra distance with Ward’s D2 linkage. Columns represent individual telomere features. Row annotations indicate ALTitude prediction (ALT+ or ALT−), experimental ALT assay strength (C-circle/APB status, where available), mutation status (*ATRX* biallelic loss, *ATRX* other, *SMARCAL1*, *DAXX*, or wild-type), *TERT* status (promoter mutation, other *TERT* mutation, or wild-type), *SMARCAL1* dependency classification (strongly dependent: Chronos score ≤ −0.5, moderately dependent: Chronos score ≤ −0.35, non-dependent: Chronos score > −0.35), mutation status (*ATRX* biallelic loss, *ATRX* other, *SMARCAL1*, *DAXX*, or wild-type), *TERT* status (promoter mutation, other *TERT* mutation, or wild-type), and age category (adult, pediatric, or unknown).Cell lines predicted as ALT+ cluster with elevated neotelomere counts, high telomere fusion rates, and low or absent *TERT* expression.

**Extended Data Fig 6.**
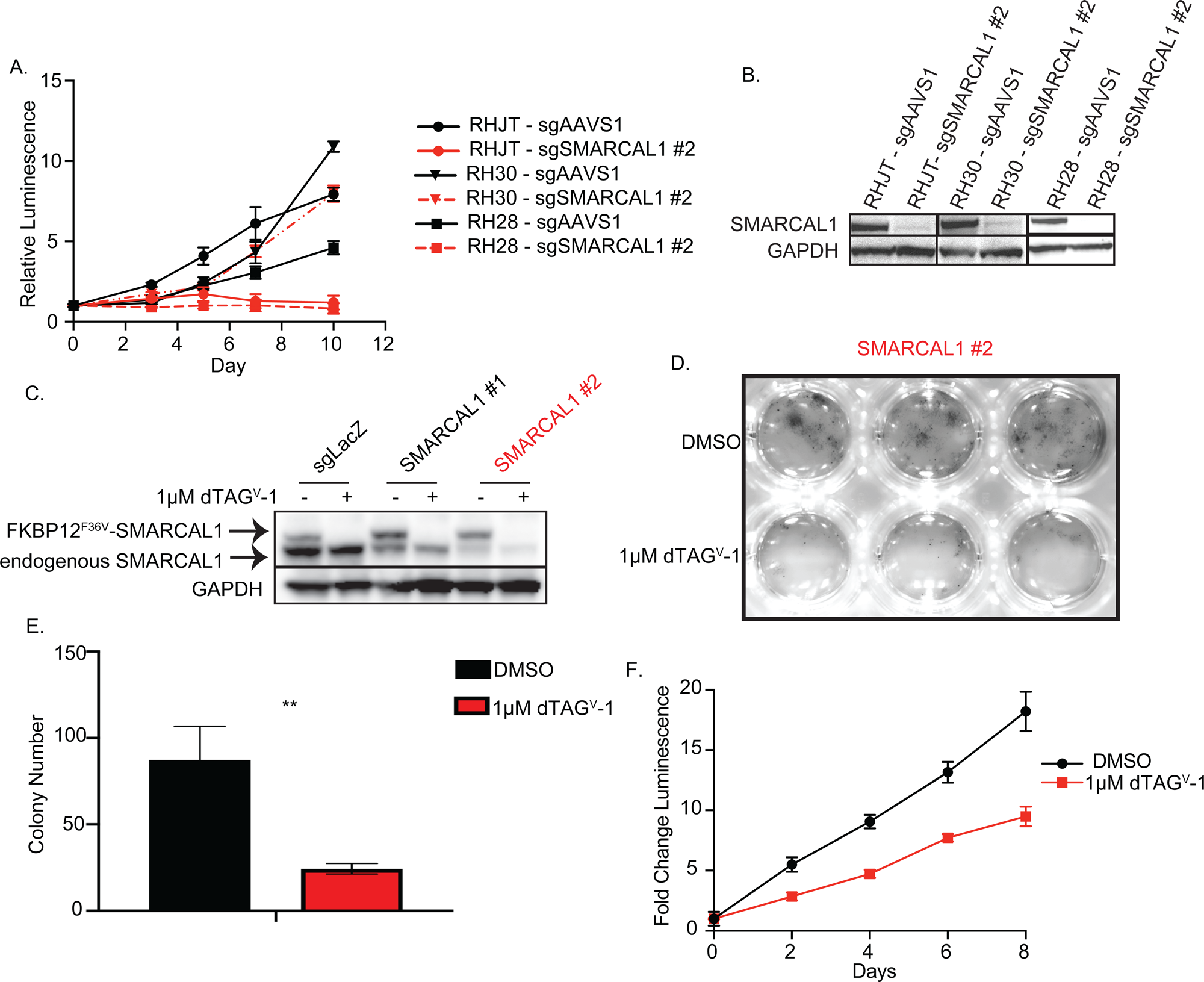
Related to Figure 2. **a.** Viability assay demonstrating *SMARCAL1* CRISPR KO (*SMARCAL1* #2) vs. gene desert control (AAVS1) in ALT+ rhabdomyosarcoma cell lines RHJT, RH30, and RH28. X-axis represents time point of luminescence measurement; y-axis represents the fold change in luminescence. **b.** Western immunoblotting confirming *SMARCAL1* KO with single guide #2 and a control (AAVS1) in the rhabdomyosarcoma lines shown in **a.** GAPDH is used as a loading control. **c.** Western immunoblotting depicting overexpression of FKBP12^F36V^-SMARCAL1 and endogenous CRISPR KO of non-targeting control (LacZ) or *SMARCAL1*, with subsequent treatment with 1µM dTAG^V^-1. GAPDH is used as a loading control. *SMARCAL1* #2 guide showed the best endogenous KO, so was chosen for methylcellulose assay. **d.** Representative images of colony formation assay shown in **c. e.** Bar graph depicting quantification (triplicate wells) of number of colonies in methylcellulose 12 days after plating with DMSO or 1µM dTAG^V^-1. **p=0.005 by unpaired T-test. **f.** Viability assay demonstrating differential growth of FKBP12^F36V^ c-terminal tagged SMARCAL1 expressing G292 cells treated with 1 µM dTAG^V^-1 or DMSO, with CRISPR KO of endogenous *SMARCAL1* (#2 in **c.**). X-axis represents time point of luminescence measurement; y-axis represents the fold change in luminescence.

**Extended Data Figure 7:**
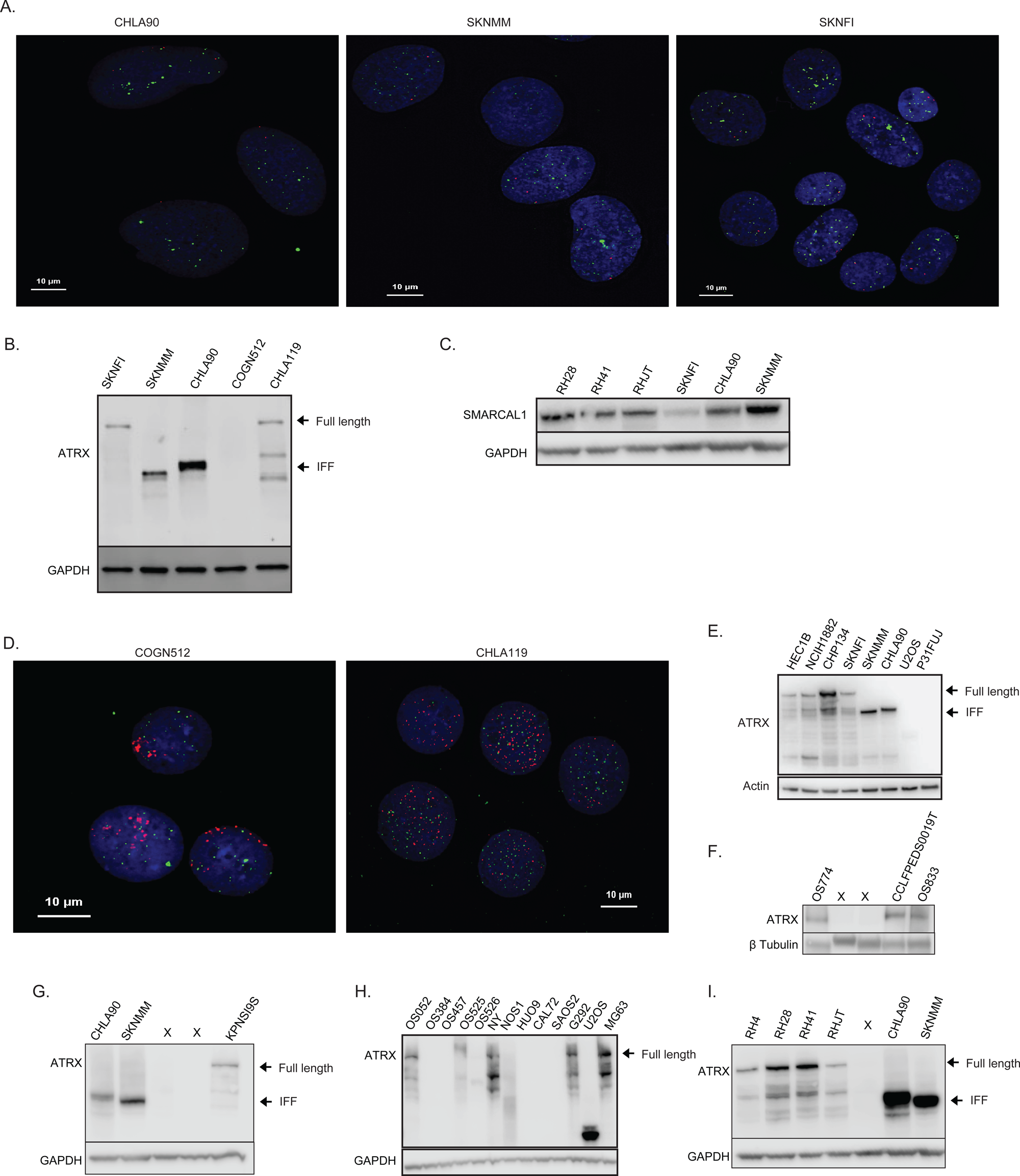
Related to Figure 3. **a.** Fluorescence in situ hybridization (FISH) performed with probes against telomere repeat sequence TTAGGG (green) and NMYC (red) in ALT+ neuroblastoma cell lines SKNFI, SKNMM, and CHLA90. Blue represents DAPI (nuclei). Scale bar 10 µm. **b.** Western immunoblot depicting different *ATRX* variants in neuroblastoma cell lines (WT vs. IFF), SKNFI, SKNMM, CHLA90, COGN512, and CHLA119. GAPDH is used as a loading control. **c.** Western immunoblot depicting SMARCAL1 levels in neuroblastoma ALT+ cell lines (SKNFI, CHLA90, SKNMM) and rhabdomyosarcoma cell lines (RH28, RH41, RHJT). GAPDH is used as a loading control. **d.** Fluorescence in situ hybridization (FISH) performed with probes against telomere repeat sequence TTAGGG (green) and NMYC (red) in neuroblastoma cell lines COGN512 (ALT+) and CHLA119 (non-ALT). Blue represents DAPI (nuclei). Scale bar 10 µm. **e-i.** Western immunoblots showing ATRX size/absence in a panel of cell lines to confirm mutation characteristics assessed by WGS. GAPDH, β-Tubulin, or Actin are used as loading controls. All used antibodies probing the C-terminus of ATRX (**e.-f.** Sigma HPA001906, **g.-i.** Cell Signaling 10321). In **e.**, **g., and i.** CHLA90 and SKNMM (with ATRX-IFF) are shown for size reference compared with full-length ATRX.

**Extended Data Figure 8:**
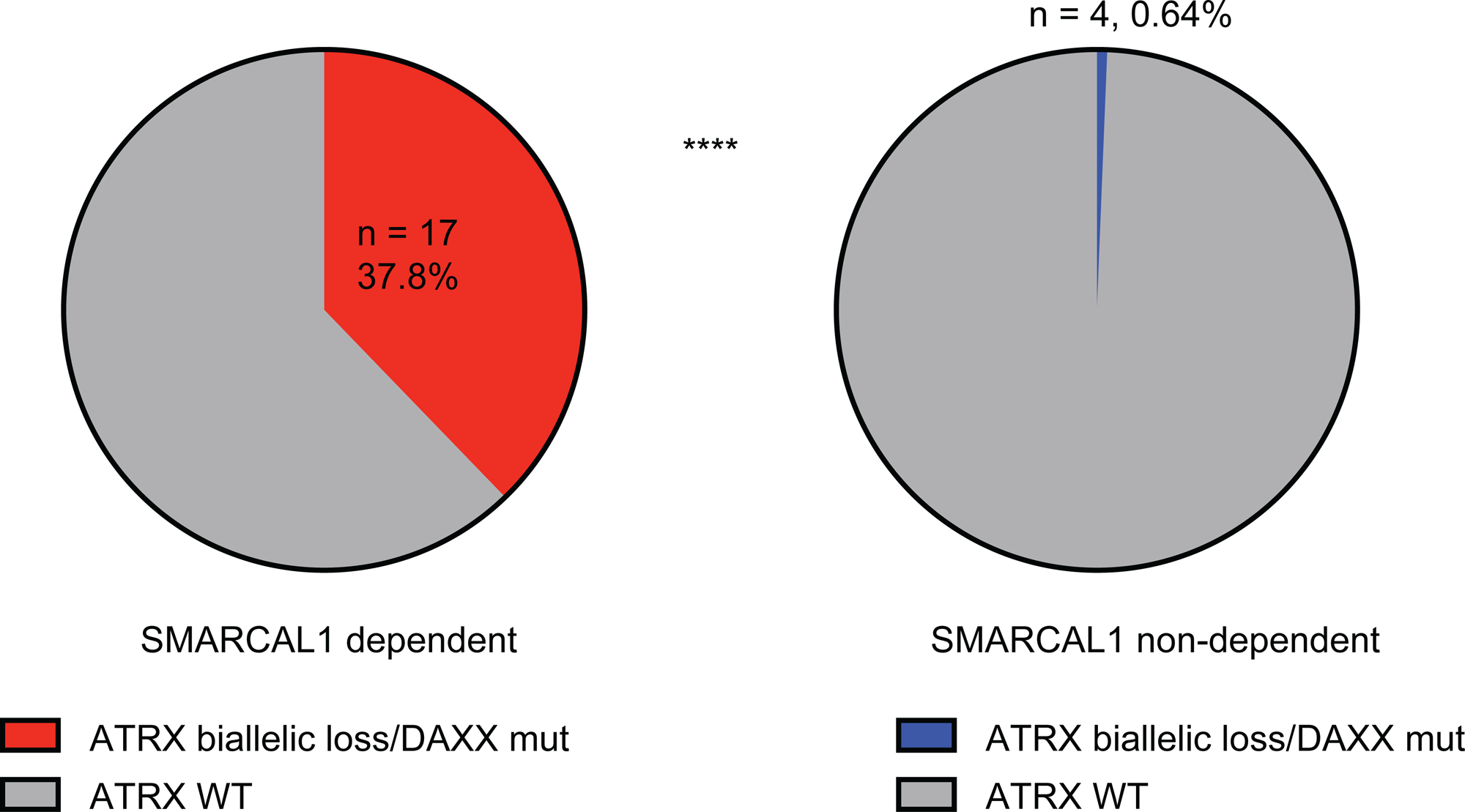
Related to Figure 4. Pie chart demonstrating proportions of *SMARCAL1* dependent vs. non-dependent cell line models with *ATRX* biallelic loss or *DAXX* mutation vs. WT. **** p<0.0001 by Fisher’s Exact test.

**Extended Data Figure 9:**
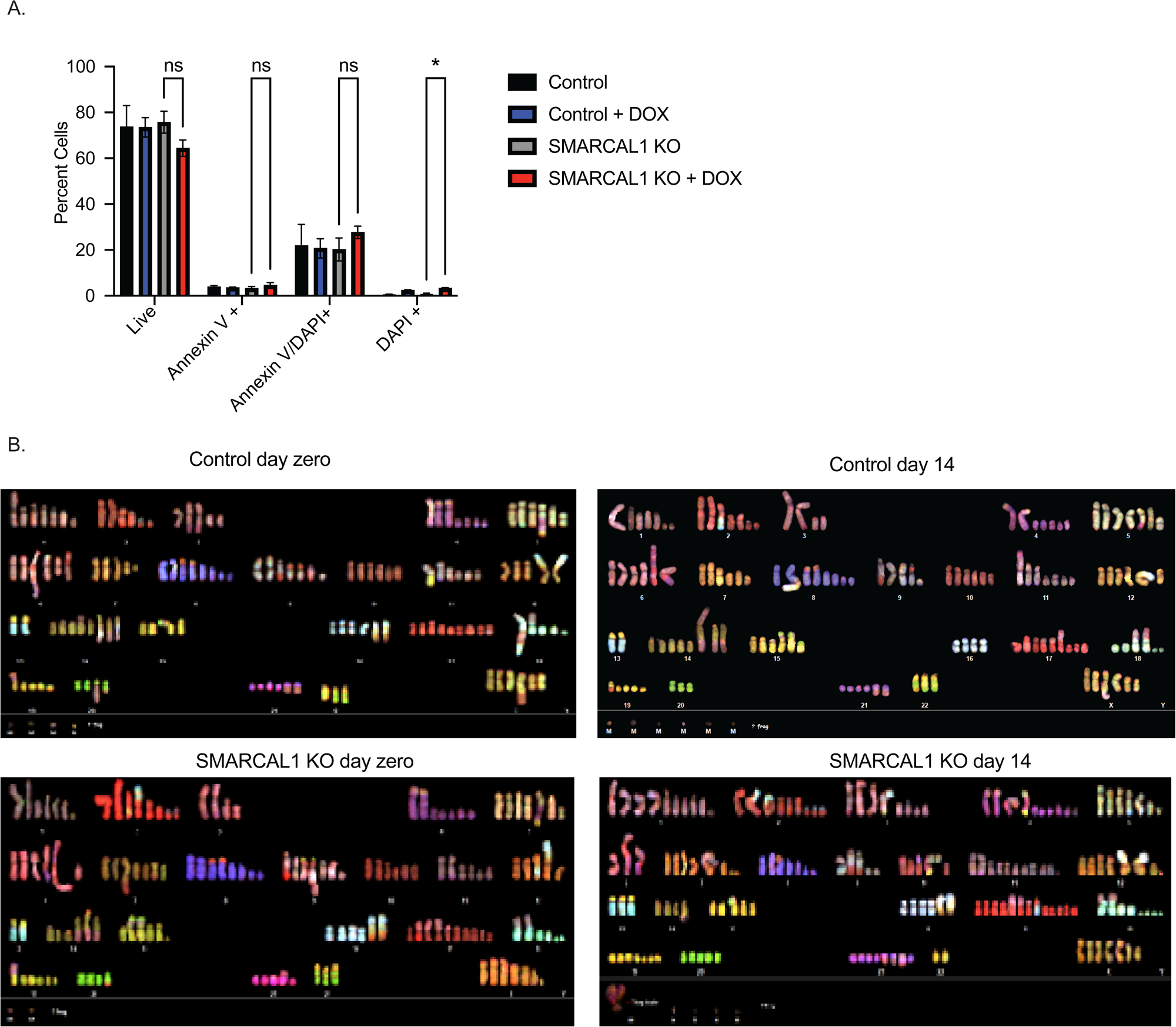
Related to Figure 6. **a.** Bar graphs demonstrating annexin/PI apoptosis flow cytometry cell counts at fourteen days post-doxycycline induction of CRISPRi knockdown of *SMARCAL1* vs. gene desert control (Chr2.2.) vs. non-doxycycline induced controls. Proportions of live, Annexin +, Annexin+ and PI+, cells are shown. Only dead cells are significant *p<0.05 by unpaired T-test. **b.** SKY sample images demonstrating number of chromosomes/fragments on day zero (3 days post dox-selection) and day 14 following gene desert control (Chr2.2) KO or *SMARCAL1* KO.

## Notes

### Competing Interest Statement

The authors have declared no competing interest.

### Summary of Updates

Updated Data, figures revised & text revised.

